# Cellular and Synaptic Phenotype Compensations Limit Circuit Disruption in *Fmr1-KO* Mouse Layer 4 Barrel Cortex but Fail to Prevent Deficits in Information Processing

**DOI:** 10.1101/403725

**Authors:** Aleksander P.F. Domanski, Sam A. Booker, David J.A. Wyllie, John T.R. Isaac, Peter C. Kind

**Affiliations:** School of Physiology, Pharmacology & Neuroscience, University of Bristol, Bristol, UK; Centre for Discovery Brain Sciences, University of Edinburgh, Hugh Robson Building, George Square, Edinburgh, EH8 9XD, UK; Patrick Wild Centre, University of Edinburgh, Hugh Robson Building, George Square, Edinburgh, EH8 9XD, UK; Simons Initiative for the Developing Brain, University of Edinburgh, Hugh Robson Building, George Square, Edinburgh, EH8 9XD, UK; Centre for Brain Development and Repair, NCBS, GKVK Campus, Bangalore, 560065, India; Developmental Synaptic Plasticity Section, NINDS, NIH, Bethesda, MD 20892, USA

## Abstract

Sensory hypersensitivity is a common and debilitating feature of neurodevelopmental disorders such as Fragile X Syndrome (FXS). However, how developmental changes in neuronal function ultimately culminate in the network dysfunction that underlies sensory hypersensitivities is not known. To address this, we studied the layer 4 barrel cortex circuit in *Fmr1* knockout mice, a critical sensory processing circuit in this mouse model of FXS. By systematically studying cellular and synaptic properties of layer 4 neurons and combining with cellular and network simulations, we explored how the array of phenotypes in *Fmr1* knockout produce circuit pathology during development that result in sensory processing dysfunction. We show that many of the cellular and synaptic pathologies in *Fmr1* knockout mice are antagonistic, mitigating circuit dysfunction, and hence can be regarded as compensatory to the primary pathology. Despite this compensation, the layer 4 network in the *Fmr1* knockout exhibits significant alterations in spike output in response to ascending thalamocortical input that we show results in impaired sensory encoding. We suggest that it is this developmental loss of layer 4 sensory encoding precision that drives subsequent developmental alterations in layer 4 – layer 2/3 connectivity and plasticity observed in the *Fmr1* knockout, and is a critical process producing sensory hypersensitivity.

## Introduction

Individuals affected by many types of Autism Spectrum Disorder (ASD) and Intellectual Disabilities (ID) commonly exhibit sensory perceptual disturbances that span multiple modalities (Rogers, Hepburn and Wehner, 2003; Crane, Goddard and Pring, 2009; Marco *et al.*, 2011). Fragile X Syndrome (FXS) is the leading heritable cause of ASD/ID (Hagerman *et al.*, 2009) with symptoms including seizures, tactile hypersensitivity and abnormal behaviours that affect early sensory and cognitive development. FXS is caused by loss of FMRP protein following transcriptional silencing of the *Fmr1* gene. Like FXS, the *Fmr1-KO* mouse model (Consortium, 1994) lacks FMRP and exhibits sensory, behavioural and cognitive deficits (Bernardet and Crusio, 2006; Mineur, Huynh and Crusio, 2006). Sensory dysfunction in FXS and related ASDs have been proposed to underlie a range of behavioural and cognitive symptoms. In support of this hypothesis, a recent study has indicated a causal link between sensory dysfunction and social and repetitive behaviours in a mouse model of autism (Shin Yim *et al.*, 2017). Hence a detailed understanding the sensory function in FXS may be critical to developing novel therapies.

Rodent models demonstrate that the sensory hypersensitivities associated with *Fmr1* deletion are mirrored by an increase in circuit excitability (Zhang *et al.*, 2012; Gonçalves, Anstey, Golshani and Portera-Cailliau, 2013; He *et al.*, 2017, 2018; O’Donnell *et al.*, 2017). However, the known cellular processes that contribute to circuit hyperexcitability in *Fmr1-KO* mice are numerous (Contractor, Klyachko and Portera-Cailliau, 2015) and the potential number is even greater. FMRP has the potential to regulate the translation of diverse classes of neuronal mRNA (Brown et al., 2001; Darnell et al., 2001, 2011) including ion channels, neurotransmitter receptor subunits, and intracellular signalling molecules. Furthermore, protein-protein interactions with voltage gated ion channels directly link FMRP to maintenance of intrinsic neuronal properties (Brown *et al.*, 2010; Strumbos *et al.*, 2010; Zhang *et al.*, 2012; Deng *et al.*, 2013). Finally alterations of the Inhibitory/Excitatory (In/Ex) balance have been reported in *Fmr1-KO* mice (Gibson *et al.*, 2008; Paluszkiewicz *et al.*, 2011; Gonçalves, Anstey, Golshani and Portera-cailliau, 2013; Vislay *et al.*, 2013; Cea-Del Rio and Huntsman, 2014; He *et al.*, 2014) although the precise cellular mechanism underlying this In/Ex imbalance is unknown. Hence, a detailed dissection of the contribution of individual cellular phenotypes underlying the emergent circuit pathophysiology is required to understand the sensory processing deficits associated with FXS (O’Donnell *et al.*, 2017).

In the somatosensory cortex of *Fmr1-KO* mice, circuit dysfunction arises very early in development correlating with the peak of FMRP expression during the second postnatal week (Harlow *et al.*, 2010), a developmental stage marked by both the end of critical period refinement for thalamocortical (TC) synaptic input and the coordinated maturation of cortical layer 4 cell-intrinsic properties and recurrent network circuitry (Daw, Bannister and Isaac, 2006; Daw, Ashby and Isaac, 2007; Daw, Scott and Isaac, 2007; Chittajallu and Isaac, 2010; Ashby and Isaac, 2011). Loss of FMRP delays the onset and termination of the critical period for synaptic plasticity at TC synapses (Harlow et al., 2010). The consequences of this delay, both in terms of cellular and circuit function, are unknown, however, it is notable that active whisking begins soon after (Landers and Philip Zeigler, 2006). Hence, changes in the cellular and circuit physiology in layer 4 at this age could dramatically alter the nature of the sensory information being transmitted to layer 2/3 that drives further experience-dependent development. Importantly, at later ages *Fmr1*-*KO* mice display disrupted functional connectivity in layer 4 and an altered synaptic In/Ex balance (Gibson et al., 2008; Paluszkiewicz et al., 2011b; Baudouin et al., 2012; Cellot and Cherubini, 2014). However, whether these differences arise as a direct result of the loss of FMRP or are compensatory changes resulting from earlier developmental alterations in cellular physiology is not known. Furthermore, it is unclear if the cellular abnormalities that result from a delay in the sensitive period for synaptic plasticity at TC synapses underlies the altered In/Ex balance.

To address these questions, we examined the cellular and circuit properties of excitatory and feedforward inhibition providing Fast-spiking (FS) GABAergic neurons in layer 4 of barrel cortex at postnatal days 10-11 (P10/11, Miller, Pinto and Simons, 2001; Daw, Scott and Isaac, 2007; Favorov and Kursun, 2011), immediately after the termination of the delayed sensitive period for synaptic plasticity in *Fmr1-KO* mice and immediately prior to the onset of whisking behaviour. By combining brain slice electrophysiology and computational modelling, we show that an array of cellular level pathologies is observed in *Fmr1-KO* layer 4: in connectivity, in cellular intrinsic properties and in synaptic function. The net result of these pathologies is a circuit with a lower threshold for action potential generation in response to TC input, but less well-timed firing relative to input stimuli. Our modelling shows that the cellular-level pathologies observed in the *Fmr1-KO* are often antagonistic in terms of circuit function suggesting that some ‘pathologies’ rather may be compensatory adaptations. Despite the compensation, the layer 4 circuit is dysfunctional in *Fmr1-KO* with a reduced ability to encode information manifesting as a reduction in pattern classification accuracy as relayed to layer 2/3.

## Results

### Alterations in passive membrane properties and excitability in SCs and FS in *Fmr1-KO*

Previous work has identified a number of cellular, synaptic and circuit changes in *Fmr1-KO* mouse barrel cortex (Gibson *et al.*, 2008; Hays, Huber and Gibson, 2011; Gonçalves, Anstey, Golshani and Portera-Cailliau, 2013). However, these experiments were conducted 5 days after the end of the delayed critical period for Long-term potentiation (LTP) in *Fmr1-KO* mice. Therefore, it is unclear whether these changes are pathological or compensatory to network function or indeed how these cellular and synaptic mechanisms interact to produce circuit level deficits. To address this, we systematically analysed cellular, synaptic and network changes in thalamocortical brain slices (Agmon and Connors, 1991; Feldman, Nicoll and Malenka, 1999) from younger *Fmr1-KO* mice immediately following the period of LTP at thalamocortical synapses and evaluated the effects of these mechanisms at a circuit level using computer simulations of layer 4 barrel cortex.

We first investigated whether there were changes in passive membrane properties in layer 4 barrel cortex neurons in acute slices from P10/11 *Fmr1*-*KO* mice compared to wild-type (WT) littermates, using whole-cell patch-clamp recordings. The principal cell type in layer 4 barrel cortex is the stellate cell (SC), which are recurrently connected glutamatergic neurons that project to layer 2/3 (Feldman, 2000; Feldmeyer and Sakmann, 2002; Helmstaedter *et al.*, 2008; Lefort *et al.*, 2009; Feldmeyer *et al.*, 2012). Assessing intrinsic excitability, compared to WTs, *Fmr1-KO* SCs required less injected current to fire an action potential (lower rheobase) and exhibited an increase in both input resistance and membrane time constant (Figure 1A), but no change in membrane capacitance or resting membrane potential (Figure S1). SCs in *Fmr1-KO*s also had enhanced excitability with an increase in the number of action potentials elicited by depolarising current steps (Figure 1B); however, this increase in action potential number was associated with a decrease in action potential frequency during the early part of the depolarisation resulting in a reduction in action potential frequency adaptation (Figure 1C). Moreover, in agreement with a previous finding in *Fmr1-KO* mouse hippocampus (Deng *et al.*, 2013) action potential kinetics were slowed in *Fmr1*-*KO*s, with an increase in width and a decrease in amplitude (Figure 1D).

**Figure 1.**
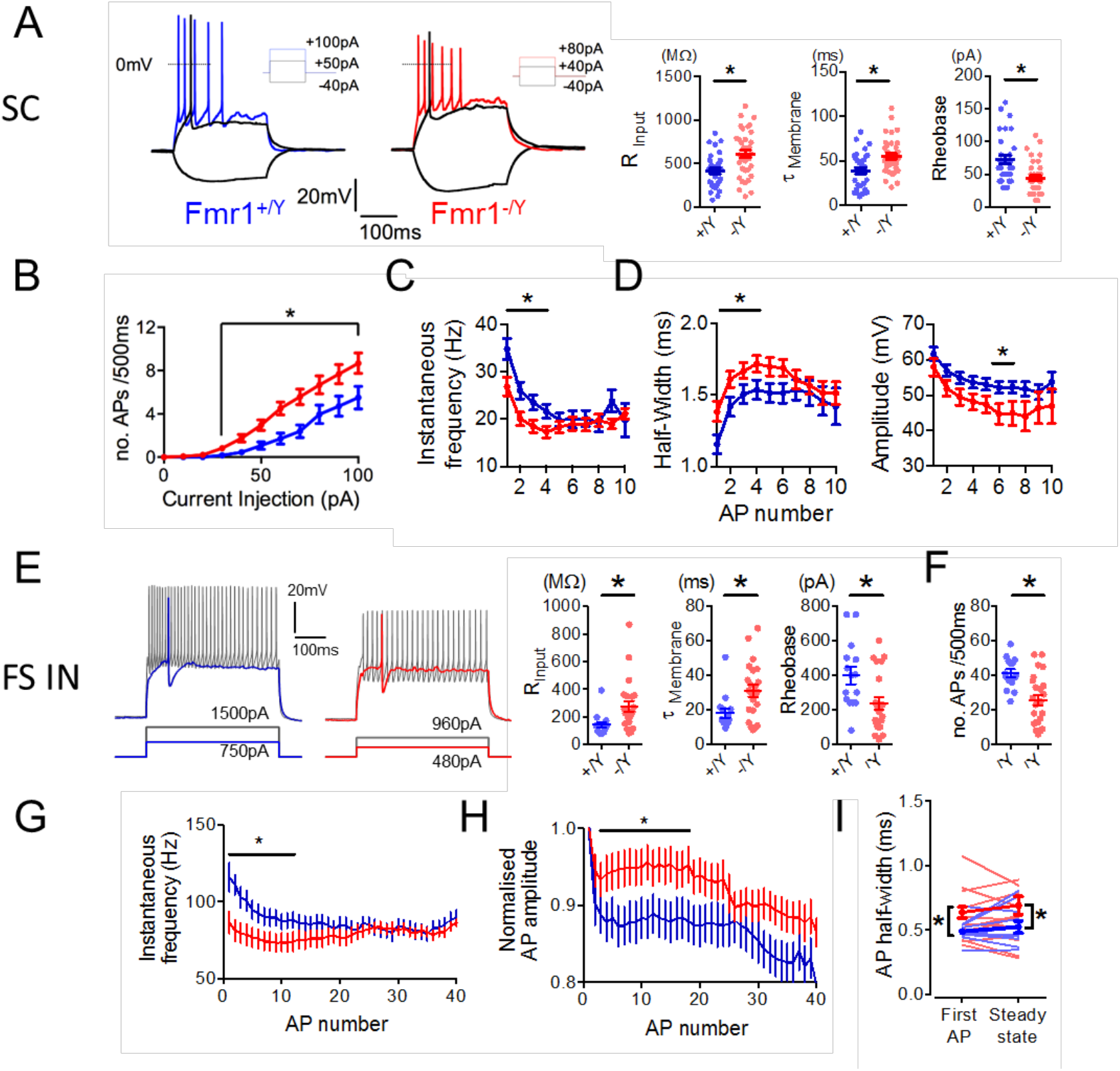
Altered intrinsic properties of *Fmr1-KO* Layer 4 SCs and FS interneurons. **A).** Left: Example firing characteristics of layer 4 excitatory neurons in response to 500ms hyperpolarizing and depolarizing current injections (-40pA, rheobase, 2x rheobase shown). Right: Passive membrane and intrinsic properties of L4 excitatory neurons: Input resistance (WT: 412±33MΩ, *Fmr1-KO*: 609±43MΩ, p=0.0007), Membrane time constant (WT: 39±3.7ms, *Fmr1-KO*: 55±3.2ms, p=0.0015), Rheobase current (P<0.002, WT: 72±6.6pA; N=33, *Fmr1-KO*: 44±4.0pA; N=37). Not shown: Membrane capacitance (WT: 94±6.7pF, *Fmr1-KO*: 89±4.9pF, p=0.56), resting membrane potential (WT: -64±1.7mV, *Fmr1-KO*: -64±1.3mV, p=0.88). **B).** Suprathreshold current-spike frequency (FI) responses of L4 *Fmr1-KO* excitatory neurons were significantly steeper during 500ms depolarizing current injections >30pA (p=0.02, Mann-Whitney, WT: 120±10Hz/nA, *Fmr1-KO:* 200±12Hz/nA, (n: 28 WT, 28 *Fmr1-KO*). **C).** SC action potential rate during entrained firing to twice-rheobase current injections. Asterisks indicate significantly different parameters (p<0.05, t-tests comparing values for each spike position in train, N: WT=50 neurons, *Fmr1-KO* =42 neurons). **D).** SC action potential half-width and amplitude during entrained firing to twice-rheobase current injections. Asterisks indicate significantly different parameters (p<0.05, t-tests comparing values for each spike position in train, N: WT=50 neurons, *Fmr1-KO* =42 neurons). **E).** Left: Example spike waveforms fired by FS interneurons in response to 500ms depolarizing current injections (rheobase, 2x rheobase shown). Right passive membrane and intrinsic properties of FS interneurons (Asterisks indicate p<0.05, t-test, N (neurons): WT=15, Fmr1-KO =23). **F).** FS overall action potential rate during entrained firing to twice-rheobase current injections. Statistics as **E).** **G).** Accommodation of FS action potential rate during entrained firing to twice-rheobase current injections (Asterisks indicate p<0.05, t-test, N (neurons): WT=15, Fmr1-KO =23). **H).** Accommodation of FS action potential amplitude during entrained firing to twice-rheobase current injections (Asterisks indicate p<0.05, t-test, N (neurons): WT=15, Fmr1-KO=23). **I).** Reduced rate of AP firing in *Fmr1-KO* FS interneurons as analysed by instantaneous frequency (1/inter-spike interval) of first and last two APs in train Light colours indicate individual neurons, thick bars/lines are mean±SEM for each genotype. Asterisks on left and right graphs indicate parameters significantly different between genotypes compared by t-test (p<0.05, N (neurons): WT=15, *Fmr1-KO* =27). Asterisks on centre graph indicate significantly different mean frequencies between genotypes compared by one-way ANOVA with Bonferroni’s correction for multiple comparisons (p<0.05, N: WT =15, *Fmr1-KO* =27).

A similar analysis was performed on the intrinsic excitability of fast-spiking (FS) interneurons. FS provide strong feed-forward inhibition onto SCs and play a critical role in determining the integration of TC input and action potential output of SCs (Miller, Pinto and Simons, 2001; Swadlow, 2002; Gabernet *et al.*, 2005; Daw, Bannister and Isaac, 2006; Daw, Ashby and Isaac, 2007). Similar to SCs, we observed an increase in input resistance and membrane time constant and reduced rheobase (Figure 1E), and no change in whole-cell capacitance or resting membrane potential (Figure S1B). However in contrast to SCs, FS exhibited a reduction in action potential number during depolarisation (Figure 1F) accompanied by a decrease in both action potential frequency and frequency accommodation (Figure 1 G) and a slowing of action potential kinetics (Figure 1 H,I).

Thus, in layer 4 of P10/11 *Fmr1-KO* mice both SCs and FS have altered passive membrane properties such that they produce action potentials in response to less depolarisation, compared to wild type. During sustained depolarisation, *Fmr*1*-KO* SCs continue to exhibit increased action potential output albeit with alterations in frequency and kinetics. In contrast, FS show decreased action potential output in response to sustained depolarisation as well as changes in frequency and kinetics.

### Increased response to low frequency membrane depolarisations in SCs in *Fmr1-KO*

SCs are the output neurons from layer 4 and as such play an important role to integrate ascending TC input and provide output to layer 2/3, encoding stimulus features such as the presence of an object and its surface quality in spike frequency and times (Miller, Pinto and Simons, 2001; Favorov and Kursun, 2011). The capacity of neurons to faithfully follow rhythmic synaptic input – impedance – is fundamentally constrained by their intrinsic properties (Koch, 1999). We therefore next investigated whether the changes in intrinsic properties of SCs in *Fmr1-KO* mice alter their ability to transform inputs at ethologically relevant frequencies associated with somatosensation (Ferezou *et al.*, 2007; Jadhav, Wolfe and Feldman, 2009; O’Connor *et al.*, 2010; Crochet *et al.*, 2011; Xu *et al.*, 2012). We assessed SC impedance properties (Erchova *et al.*, 2004; Lawrence, Statland, *et al.*, 2006), using a sinusoidal current of progressively increasing frequency and measured the resulting voltage response. SCs from *Fmr1-KO* mice exhibited an increased impedance between 0.5 – 4 Hz (Figure S2A), consistent with predictions from the change in passive membrane properties (Figure S3). This frequency-dependent response suggests that SCs in *Fmr1-KO* mice should exhibit alterations in action potential generation in response to membrane depolarisations of the same frequency range. To test this, we applied suprathreshold sinusoidal depolarisations at specific frequencies and found an increase in the number of action potentials elicited at low frequencies in *Fmr1-KO* compared to wild type mice (0.5 – 4 Hz; Figure S2B). Moreover, a significant phase shift in the timing of action potential generation was also observed, for example at 10 Hz (Figure S2C).

Thus, the alteration in the passive membrane properties of SCs in *Fmr1-KO* mice results in a selective increase in membrane depolarisation in response to low frequency depolarizing currents. This produces both an increase in action potential output and a shift in action potential timing in response to low frequency depolarising currents.

### Altered feedforward inhibition on SCs in *Fmr1-KO* mice

Feed-forward inhibition (FFI) is a critical mechanism that governs the integration of TC input by SCs (Gabernet *et al.*, 2005; Daw, Bannister and Isaac, 2006; Daw, Ashby and Isaac, 2007; Chittajallu and Isaac, 2010; Cruikshank *et al.*, 2010). In layer 4 barrel cortex FFI is mediated by FS neurons which provide the TC-evoked inhibition onto SCs, setting an integration window for SC action potential output between the onset of excitatory and the strong but delayed inhibitory inputs (Gabernet *et al.*, 2005). To determine if there are alterations in FFI in *Fmr1-KO* mice, we first investigated connectivity between FS and SCs using simultaneous whole-cell patch-clamp recordings. Similar to findings by Gibson *et al.* at P14, at P10/11 we found reductions in connectivity from SC to FS, but additionally from FS to SC and reciprocally between SC and FS, in *Fmr1-KO* mice compared to wild types (Figure 2A). In contrast, however, the strength of the connections between FS and SCs was unchanged in *Fmr1-KO* mice compared to wild types (Figure 2B) and the kinetics of the IPSCs onto SCs and of EPSCs into FS were also unchanged (Figure 2C). Furthermore, there was no difference in SC-SC connectivity in *Fmr1-KO* mice compared to wild-type, nor in synaptic strength or kinetics (Figure 2D-F).

**Figure 2.**
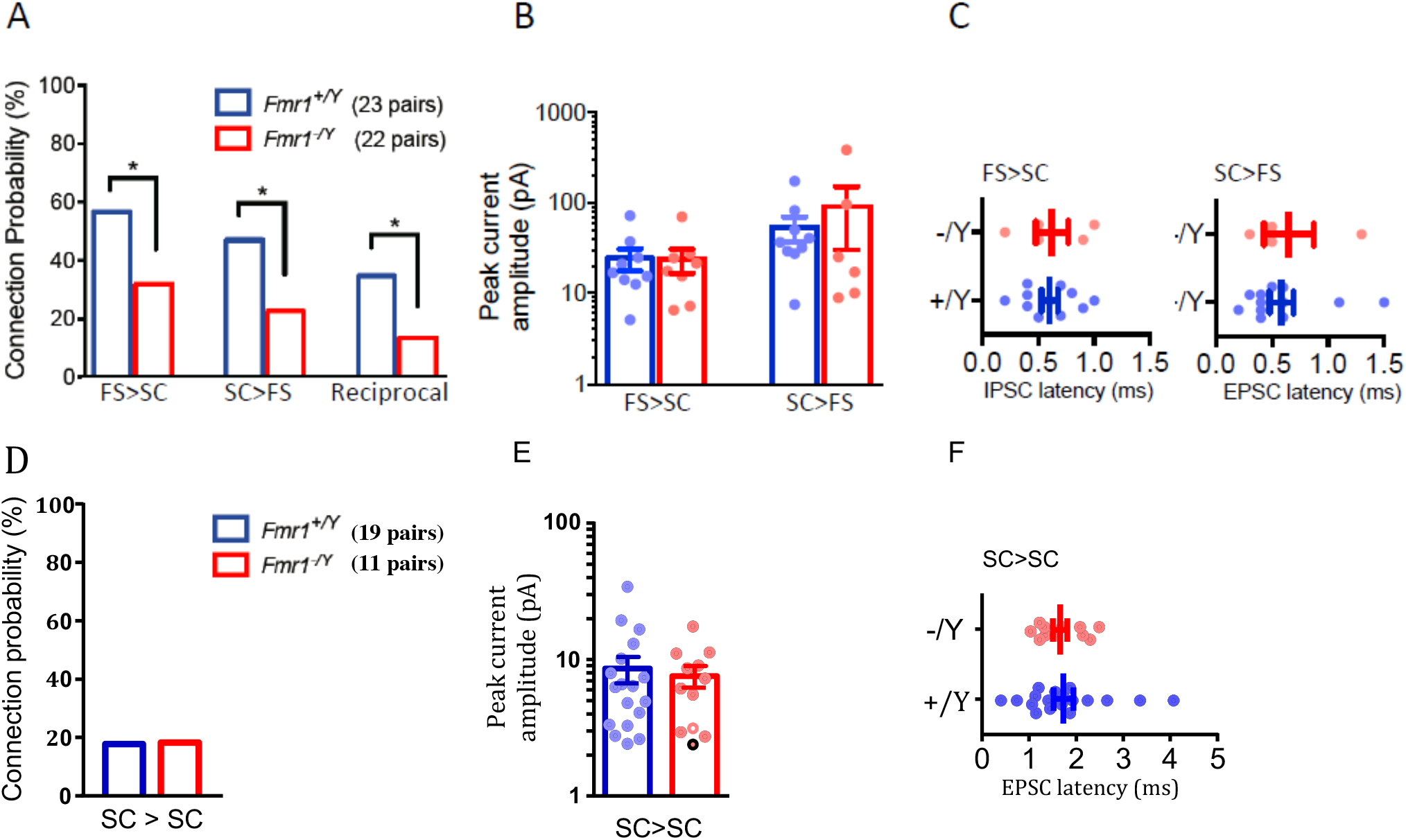
Reduced FS IN – SC connectivity in *Fmr1-KO*, but no change in connection strength or kinetics of currents. **A).** Monosynaptic connection probability between pairs of FS and SCs tested with paired whole-cell recording. Asterisks indicate statistically significant differences (p<0.05, Chi-squared test). **B-C).** Connection strengths (**B)**, peak evoked monosynaptic current amplitude and current onset latencies (**C)** for connected pairs shown in **A).** No differences in connection strength between genotype were observed for either connection direction (p>0.05, Mann-Whitney, N’s as **A)**). **D).** Monosynaptic connection probability between pairs of SCs located within the same barrel for recordings in slices taken from P10-11 (left) and P12-15 (right) mice of each genotype. No change in connection probability was observed at P10-11 but a deficit emerged by P15. Asterisk indicates p<0.05, Chi-squared test. **E).** No change in synaptic strengths between connected SC pairs shown in P10-11 *Fmr1-KO* mice shown in **D).** **F)** No change in latency to peak monosynaptic EPSC amplitude for connected pairs show in in **D).**

We next directly measured FFI in SCs, by stimulating TC inputs and comparing the size of the monosynaptic TC-evoked EPSC (at -70 mV holding potential) and the size of the di-synaptic feed forward IPSC in the same SCs (at 0 mV; Chittajallu & Isaac 2010). In SCs from *Fmr1-KO* mice, we found an increase in the peak amplitude of the feedforward IPSC relative to the monosynaptic EPSC (‘GABA/AMPA ratio’ [G/A]; Figure 3A). Notably however, a subset of *Fmr1-KO* neurons lacked any FFI, whereas FFI was observed in all tested WT neurons. The feedforward IPSC also exhibited slower kinetics and a longer onset lag time (compared to the monosynaptic EPSC) in SCs from *Fmr1-KO* mice (Figure 3B).

**Figure 3.**
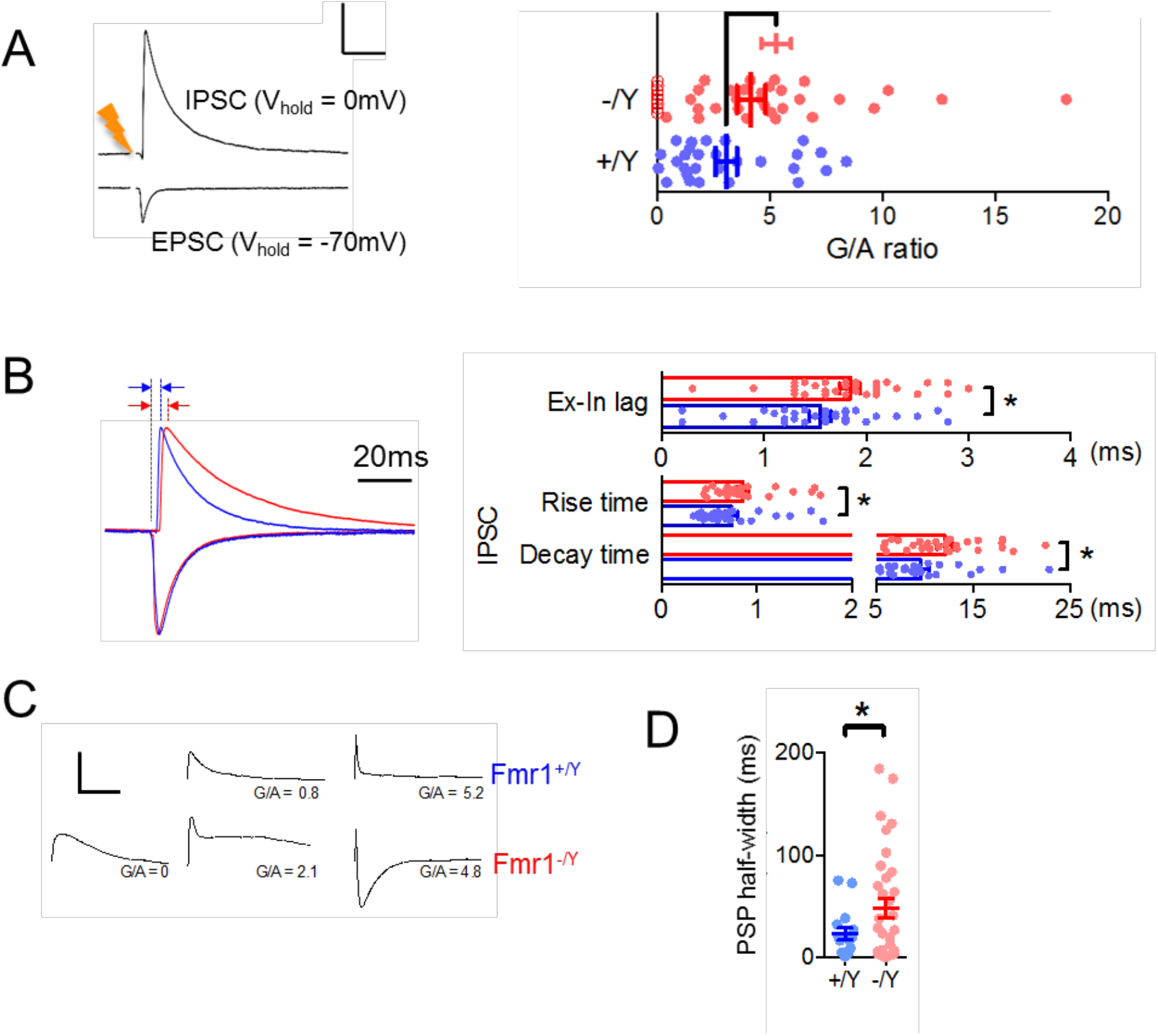
Altered thalamocortical feed-forward inhibition in P10-11 Fmr1-KO mouse. **A). Left:** Example trial-averaged (15 sweeps each) voltage-clamped whole-cell recording from a WT Layer 4 excitatory neuron, direct thalamocortical EPSCs (1) and FF-IPSCs (2) can be isolated by voltage clamping voltage clamped by holding the neuron at E_GABAa_ (-70mV) and ENMDA/AMPA (0mV), respectively. The strength of TC FFI (“G/A ratio” is quantified as the ratio of peak evoked current. Scale: 25ms/500pA. **Right:** Strength of TC-FFI at P10-11. Points indicate neurons, bars are mean±SEM. Unlike for Fmr1+/Y recordings (28 neurons, maximum of 3 neurons from each animal), in Fmr1-/Y neurons, some cells (8/38 neurons, maximum of 3 from each animal) were found to have no detectable FFI; light red bars indicate the mean±SEM strength of the remaining population of neurons (i.e. cells with G/A>0, hollow markers). Including Fmr1-/Y neurons with G/A=0, average strength of FFI was not significantly different to that of Fmr1+/Y, but excluding these neurons, (dark red bars and solid markers), the average strength was elevated over that of FFI in wild-type recordings (Fmr1+/Y vs. all Fmr1-/Y neurons: p=0.40, Fmr1+/Y vs. Fmr1-/Y neurons with G/A>0: p=0.005, Mann-Whitney). **B).** Synaptic kinetics of currents underlying TC-FFI. **Left:** Example TC-evoked currents in voltage-clamp recordings from Fmr1+/Y (blue) and Fmr1-/Y (red), EPSCs and FF-IPSCs are shown individually scaled to peak amplitudes. Note the pronounced increase in decay time constant and onset latency for Fmr1-/Y FFI-PSCs (indicated by red and blue arrows). **Right:** Slower synaptic kinetics of feed-forward inhibitory currents in recordings from *Fmr1-KO* Layer 4 excitatory neurons. No significant differences between genotypes were observed in the same comparisons for kinetics of EPSCs. Asterisks indicate significant differences (p<0.05, t-test, N’s: EPSCs N=23 (Fmr1+/Y) and N= 26 (Fmr1-/Y) neurons, FF-IPSCs N=27 (Fmr1+/Y) and N= 28 (Fmr1-/Y) neurons). **C).** Example thalamocortical EPSPs recorded from Layer 4 excitatory neurons with low, medium and high strength TC-FFI, recorded in current clamp configuration at -65mV. Note the progressive curtailment of EPSP duration with increasing FFI strength, the lack of a WT example for G/A=0, and the exaggerated IPSP for the high strength FFI *Fmr1-KO* example. **D).** TC EPSPs were slower in *Fmr1-KOs*. Plotted points are the full-width at half-height (‘half-width’) of EPSPs from layer 4 neurons (N (neurons) = 19 Fmr1+/Y, 36 Fmr1-/Y), bars are mean±SEM, and asterisks denote significance (t-test, p<0.05). Not shown: the dependence of *Fmr1-KO* thalamocortical EPSP duration on the strength of FFI is distorted. *Fmr1-KO* Neurons with weaker/no FFI (G/A ratio <2) showed specifically broadened EPSP duration (22.4±7.30ms vs. 68.4±16.ms, Fmr1+/Y vs. Fmr1-/Y, p=0.02), whereas this effect was dampened in neurons with stronger FFI (13.2±8.70ms vs. 37.1±15.8ms, Fmr1+/Y vs. Fmr1-/Y, p =0.07).

To investigate the consequences of these changes in the relative magnitude and timing of FFI we measured the resulting postsynaptic potential (PSP, Figure 3C). As we and others have previously shown (Pouille *et al.*, 2001; Gabernet *et al.*, 2005; Chittajallu and Isaac, 2010), the postsynaptic potential (PSP) half width in SCs is strongly influenced by FFI, which truncates the PSP. We found that PSP full width at half-maximum amplitude (‘half-width’) is increased in the *Fmr1-KO* mice (Figure 3D) suggesting a decrease in functional FFI, despite the increased feedforward inhibitory synaptic input onto SCs. This reduction in functional FFI is likely due to the changes in passive membrane properties of SCs; the mechanism is further explored as described later. One additional mechanism that could contribute to the lack of increase in functional FFI in SCs is a change in chloride reversal potential in the *Fmr1-KO* mice. To test this, we used perforated patch-clamp recordings and found no difference in chloride reversal potential in SCs from the two genotypes (Figure S4).

These findings, therefore, show that there is diminished functional FFI onto SCs in *Fmr1-KO* mice despite an increase in G/A ratio to a single stimulus. Considering the critical role feed forward inhibition plays in determining action potential generation and timing in SCs (Gabernet *et al.*, 2005; Daw, Bannister and Isaac, 2006; Swadlow, 2009), this deficit is likely to have important consequences for layer 4 network function at this crucial stage of activity-dependent development.

### Enhanced short-term synaptic depression in layer 4 in *Fmr1-KO* mice

Short-term synaptic plasticity is an important mechanism for information processing in cortical networks (Abbott and Regehr, 2004; Silver, 2010) including determining the efficacy of FFI (Gabernet *et al.*, 2005; Daw, Bannister and Isaac, 2006; Daw, Ashby and Isaac, 2007; Chittajallu and Isaac, 2010). We therefore compared short-term depression of the TC excitatory input and of the feed forward TC-evoked IPSC onto SCs. Both excitatory and feed forward inhibitory transmission in the *Fmr1-KO* mice showed an increase in short-term depression during trains of five stimuli over a frequency range from 5 – 50 Hz compared to WT (Figure 4A). However, the relative increase in short-term depression was greater for the IPSC than the EPSC leading to a stronger reduction in inhibitory/excitatory ratio during trains of activity in *Fmr1-KO* compared to wild type mice (Figure 4B,C). We also investigated short-term plasticity at SC-FS and FS-SC synapses in connected simultaneous recordings. In contrast to the normal short-term plasticity observed by Gibson et al. (2008) in *Fmr1-KO* recordings at two and four weeks old, we found increased short-term depression at P10/11, both for the excitatory input onto FS from SC and for the inhibitory input from FS to SC (Figure 4D,E). In SC to SC connections, although there was a trend for a decrease in short-term depression during the train, there was no robust change observed (Figure 4F,G).

**Figure 4.**
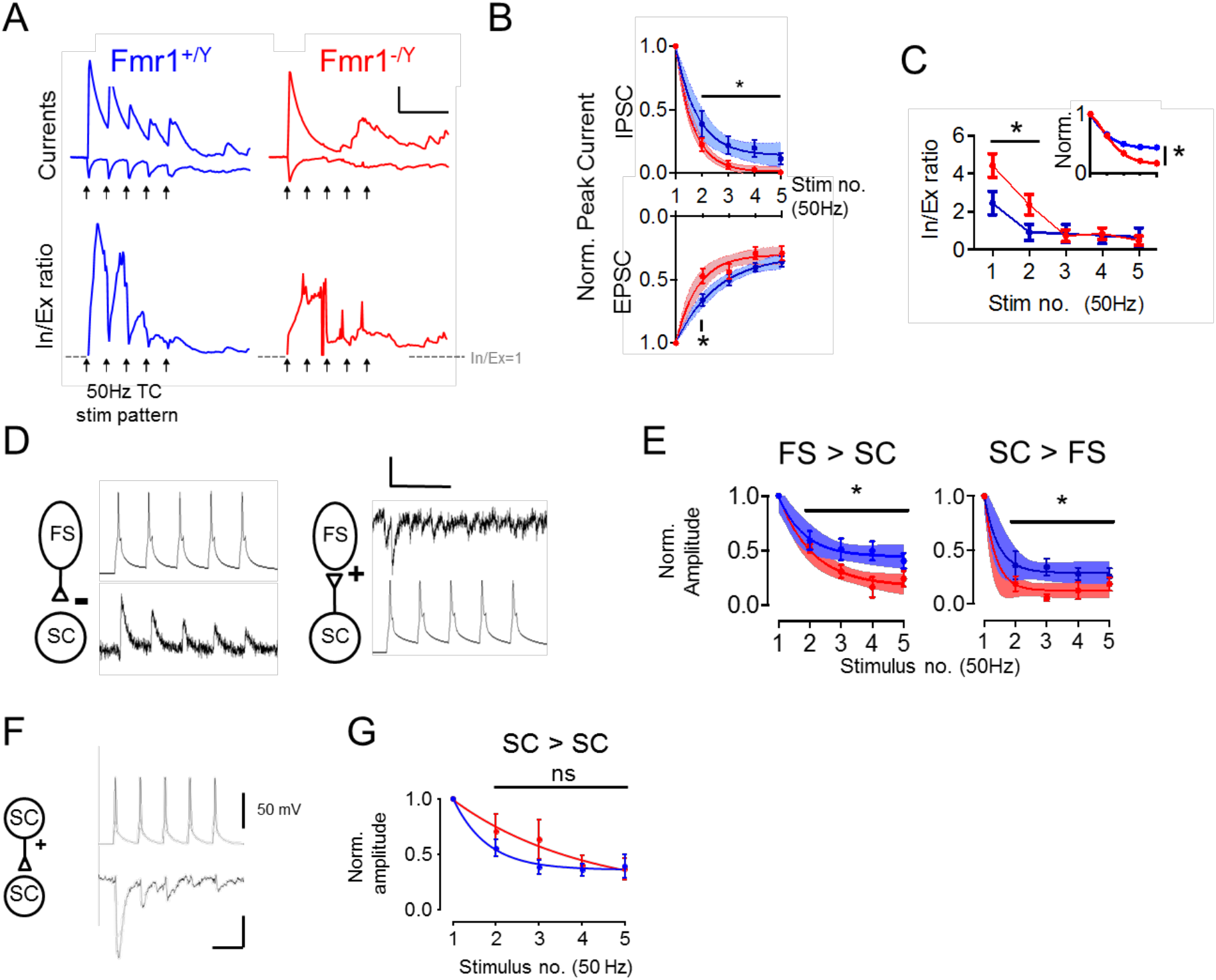
Synapse-specific changes to short-term plasticity in P10-11 *Fmr1-KO* mouse. **A).** Top: Example WT (Left) and *Fmr1-KO* (Right) voltage-clamped synaptic currents (isolated as in Figure 4A) during repetitive TC stimulation at 50Hz Traces show average of 15 trials. Note strong amplitude attenuation of TC-evoked and FFI currents in *Fmr1-KO*. Bottom: Instantaneous In/Ex ratios for the above traces calculated by dividing FF-IPSC amplitudes by EPSCs for each sampled point in time. Note: 1) smooth run-down of In/Ex ratio in the WT 2) Slow onset of In/Ex tone during stimulus train despite large amplitude FF-IPSC, 3) Temporally disorganised ratio in the *Fmr1-KO*. Arrows show stimulation times. Scale: 50ms/100pA, In/Ex = 1. **B).** Short-term depression of EPSCs and FF-IPSCs during 5x50Hz stimulation: Quantification of evoked current amplitudes as a fraction of initial (steady-state) current amplitude. Points and solid error bars indicate mean±SEM normalized current amplitude after for each stimulus for WT (blue, N=19 (EPSCs) and N=11 (FF-IPSCs) neurons) and *Fmr1-KO* (red, N=15 (EPSCs) and N=12 (FF-IPSCs) neurons). Asterisks indicate individual stimulus responses that were significantly different between genotypes (t-test, p<0.05). Shaded regions are best±95% C. I. fits to bi-exponential decay functions. For both EPSCs and FF-IPSCs, the rate of depression for *Fmr1*-KO responses was faster and a single (i.e. common) fit could not adequately explain the behaviour of both genotypes (Extra sum-of-squares F-test, EPSCs: p=0.0007, F(2,163) =7.65, IPSCs: p=0.0002, F(2,111)=9.45, N’s as above). **C).** Data from B) but converted into stimulus-by-stimulus In/Ex ratios from FF-IPSCs by EPSCs. Inset shows data normalised to starting In/Ex ratios. Asterisks indicate stimuli with significant reductions in In/Ex tone for *Fmr1-KO* data (t-test, N’s as in B).). **D).** Example short-term depression (50Hz stimulus frequency) of unitary connections between FS-SC and SC-FS neurons (Left, Right) from paired recordings. Single trials shown. Scale 50mV,10pA/50ms. **E).** Short-term plasticity analysis in B) but for unitary connections tested between connected pairs of FS and SC neurons. (Asterisks: p>0.05, t-test, fits: (p<0.05, Extra sum-of-squares F-test N: FS to SC *Fmr1^+/Y^*=9, *Fmr1-KO* =8, SC to FS: *Fmr1^+/Y^* =9, *Fmr1^-/Y^* =6). N’s indicate neurons). **F).** As for D but for example connected SC-SC paired recording. Scale 50mV,5pA/25ms. **G).** Short-term depression of SC-SC connections was indistinguishable between WT and *Fmr1-KO* P10-11 recordings (p>0.05, t-test, N = 17 *Fmr1-WT*, 11 *Fmr1-KO*).

### Increased postsynaptic summation and action potential output in SCs in *Fmr1-KO* mice

Our results thus far reveal a number of cellular and synaptic phenotypes in layer 4 barrel cortex neurons in the *Fmr1-KO* mice. SCs are the output neurons for layer 4 barrel cortex and serve to integrate TC input and transform it to action potential output, modulated by local feed forward inhibition (Swadlow, 2002; Petersen and Loos, 2007; Feldmeyer *et al.*, 2012). Therefore, to investigate the net effect of these phenotypes on the layer 4 circuit we studied the membrane potential response in SCs elicited by TC stimulation. Using whole-cell patch-clamp recordings in current clamp, we found that trains of five stimuli at frequencies from 5-50 Hz produced greater postsynaptic summation in SCs from *Fmr1-KO* mice (Figure 5A) and a larger fraction of stimuli eliciting action potentials (Figure 5B,C). We used cell-attached patch recordings to make less invasive measure of spike output. Using this approach, we confirmed that SCs from *Fmr1-KO* mice produce spike output at lower TC stimulus frequencies than wild-type. We further found that the timing precision of action potential firing was strongly impaired, exhibiting a lower instantaneous frequency and greater variability in rate and timing (Figure 6A-D, S5). These compound effects on trial-to-trial fidelity and lower firing rate by *Fmr1-KO* neurons reduced their population average spike density function (Figure 6B,C individual neurons shown in S5C,D). No significant genotype-dependent changes were observed in the number of spikes fired per trial. Thus, the net effect of the cellular and synaptic changes in the *Fmr1-KO* mice is that SCs that produce action potentials more readily, but with less precise timing.

**Figure 5.**
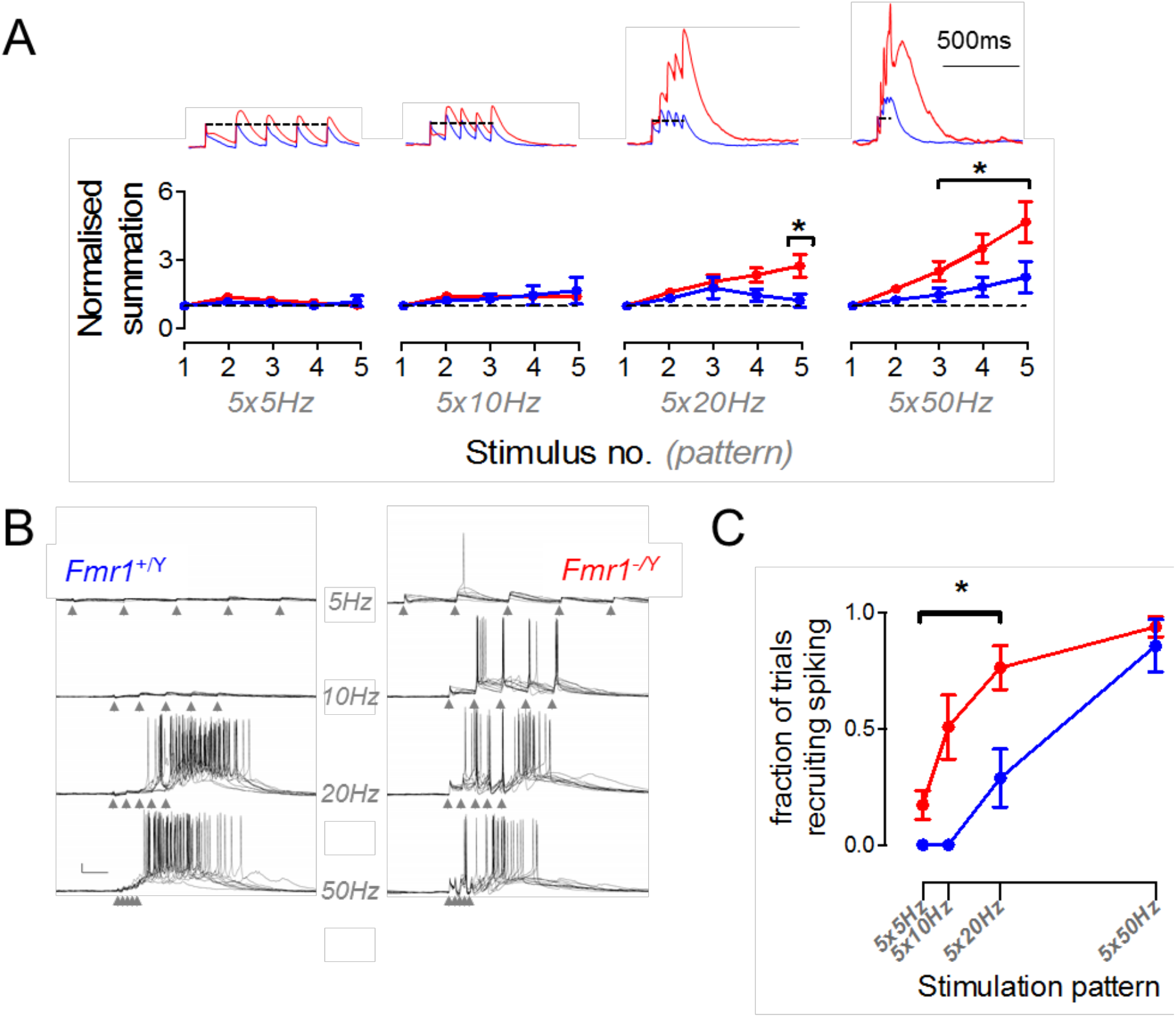
Relaxed coincidence detection of TC inputs by *Fmr1-KO* L4 excitatory neurons impairs frequency-specific gating of intracortical network activation. **A).** Hyper-summation of high-frequency thalamocortical input in *Fmr1-KOs*. Top: example current-clamp recordings showing voltage summation in response to five regular stimuli at frequencies between 5-50Hz. Amplitude is normalized to that of steady-state EPSP. Below: Normalised EPSP amplitude as a function of stimulus number in *Fmr1-KOs* and littermates. Short trains of TC stimuli at 20Hz and 50Hz evoked stronger voltage summation in *Fmr1^-/Y^* recordings: asterisks denote significantly elevated responses (P<0.05, unpaired t-test, *Fmr1^+/Y^* n=7, *Fmr1^-/Y^* n=7). **B).** Shifted sensitivity of L4 network to thalamocortical input frequency in *Fmr1-KOs*. Example current-clamp recordings (10 trials overlaid) showing transient network activity evoked by five repetitive thalamocortical stimuli at 5, 10, 20 and 50Hz. (scale bar: 100ms, 10mV). Note relaxed requirement of high-frequency stimulation for generating sustained intracortical activity in *Fmr1-KOs.* **C).** Fraction of trials evoking network activity as a function of stimulus pattern. Stimulation frequencies below 20Hz could not evoke firing (p(spiking)=0±0) in *Fmr1^+/Y^* slices, but with low-moderate probability in *Fmr1^-/Y^* slices. Asterisks denote stimulation frequencies demonstrating significantly elevated firing probabilities in *Fmr1^-/Y^* recordings (P<0.05, unpaired t-test, *Fmr1^+/Y^* n=12 slices from 8 animals, *Fmr1^-/Y^* n=10 slices from 10 animals). The 5x50Hz stimulation pattern could evoke firing with high reliability for both genotypes (p(Spiking): *Fmr1^+/Y^* = 0.87±0.11, *Fmr1^-/Y^* = 0.85±0.09).

**Figure 6.**
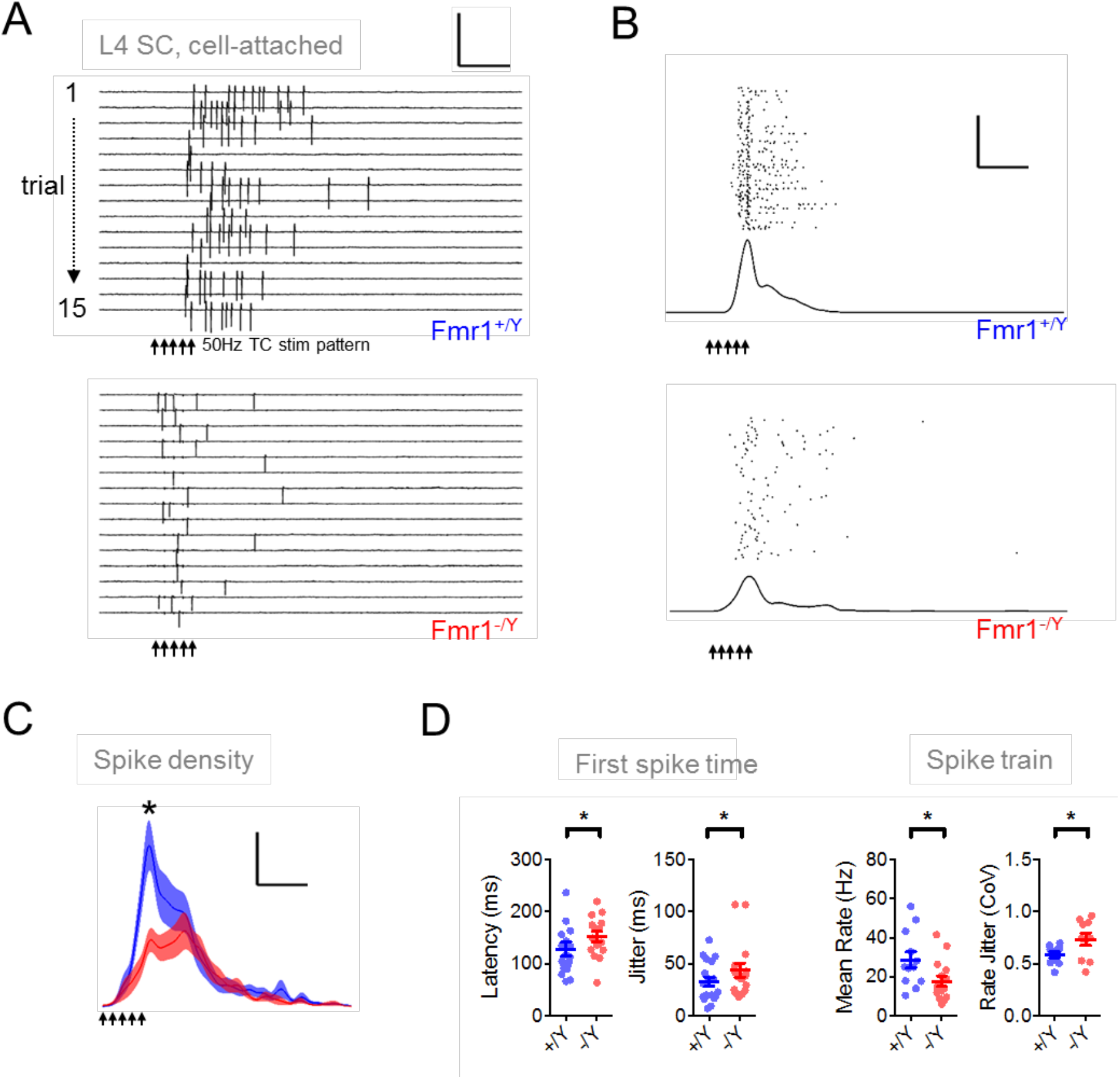
Reduced spike precision of *Fmr1-KO* Layer 4 barrel cortex excitatory neurons during thalamocortically-evoked reverberant network activity. **A).** Example multi-trial raster of spikes recorded in cell-attached configuration from WT (top) and *Fmr1-KO* (bottom) SCs in response to TC-evoked L4 network activity. Scale: 200pA/100ms. **B).** Example calculation of spike density functions for WT and *Fmr1-KO* example neurons. 200 consecutive trials (0.07Hz) showing trial-trial variability in the timing of spikes recorded in cell-attached configuration (Different cells from those shown in A).) Bottom: Trial-averaged spike density estimate (5ms kernel standard deviation) across trials for neurons shown above. Scale: 50 trials/100ms. **C).** Probability density for spikes fired by responding Layer 4 Ex neurons during the peri-stimulus period of TC-evoked network activity: mean±SEM spike density, calculated discretizing spikes to 1ms time bins and convolved with ±5ms St.Dev. Gaussian kernels. Peak mean spike density was reduced in *Fmr1-KO* recordings, calculated across the whole 1s post-stimulus sampling period (*Fmr1^+/Y^*: 0.023±0.002 sipkes.s-1, *vs. Fmr1^-/Y^*: 0.015±0.0016 spikes.s-1, p=0.008, t-test, N: *Fmr1^+/Y^*: 21 neurons, *Fmr1^-/Y^*: 16 neurons). Mean spike density averaged across the successive 200ms window of was not significantly different between genotypes (t-test, p=0.7). Scale: p(Spike/5ms) =0.5% / 100ms. **D).** Left: Spike time statistics for the first spike fired per trial for SCs. Mean latency (left) and inter-trial precision (right) were significantly slower and reduced in *Fmr1-/Y* neurons (p=0.01 and p=0.03, respectively. t-test, N’s as in C).). Right: Spike rate and rate stability was significantly different in *Fmr1-KO* recordings compared to *Fmr1-WT* littermates: SCs fired at rates that were slower and more variable between trials. Plotted points are individual neurons, bars show mean±SEM values. Asterisks indicate statistically significant differences between genotypes (p<0.05, t-test, N’s as in **C).**).

### The cellular and synaptic changes in *Fmr1-KO* mice act antagonistically in modulating layer 4 circuit function

Our experimental data show that loss of *Fmr1* produces a variety of cellular and synaptic alterations leading to an overall change in the transformation of TC input to action potential output by SCs. The various mechanisms we describe likely interact in a complex manner to produce the overall circuit phenotype. To better understand this, we first developed a single-cell model of thalamocortical integration to systematically explore the interaction of our findings in *Fmr1-KO* SCs. We used a leaky integrate-and-fire model neuron with a single dendritic segment, receiving bulk glutamatergic synaptic input connected to a soma, receiving somatic GABAergic input, representing thalamocortical and feed-forward inhibitory inputs respectively. We adjusted the intrinsic and synaptic kinetics and short-term plasticity behaviour of the model to match WT and *Fmr1-KO* mean values and altered the strength and timing of the GABAergic input (relative to that of the glutamatergic) to explore the parameter space described by our experimental data (Fig 7A). We asked which frequencies of train stimulation between 5-50Hz led to spike firing by the model (Fig 7B). In the absence of FFI (G/A = 0, GABAergic conductance silenced) both WT and *Fmr1-KO* models fired spike(s) at all tested frequencies. Gradual addition of FFI progressively restricted the ability of trains to elicit spikes to those at higher frequencies, but this effect was less pronounced in *Fmr1-KO* model compared to WT. Notably, the range of G/A over which this effect was observed are similar to the G/A ranges we measured experimentally. The *Fmr1-KO* model also reproduced the experimentally observed increases in first spike delay, spike timing jitter and spike number compared with WT (Fig 7B insert, Fig 7C).

**Figure 7.**
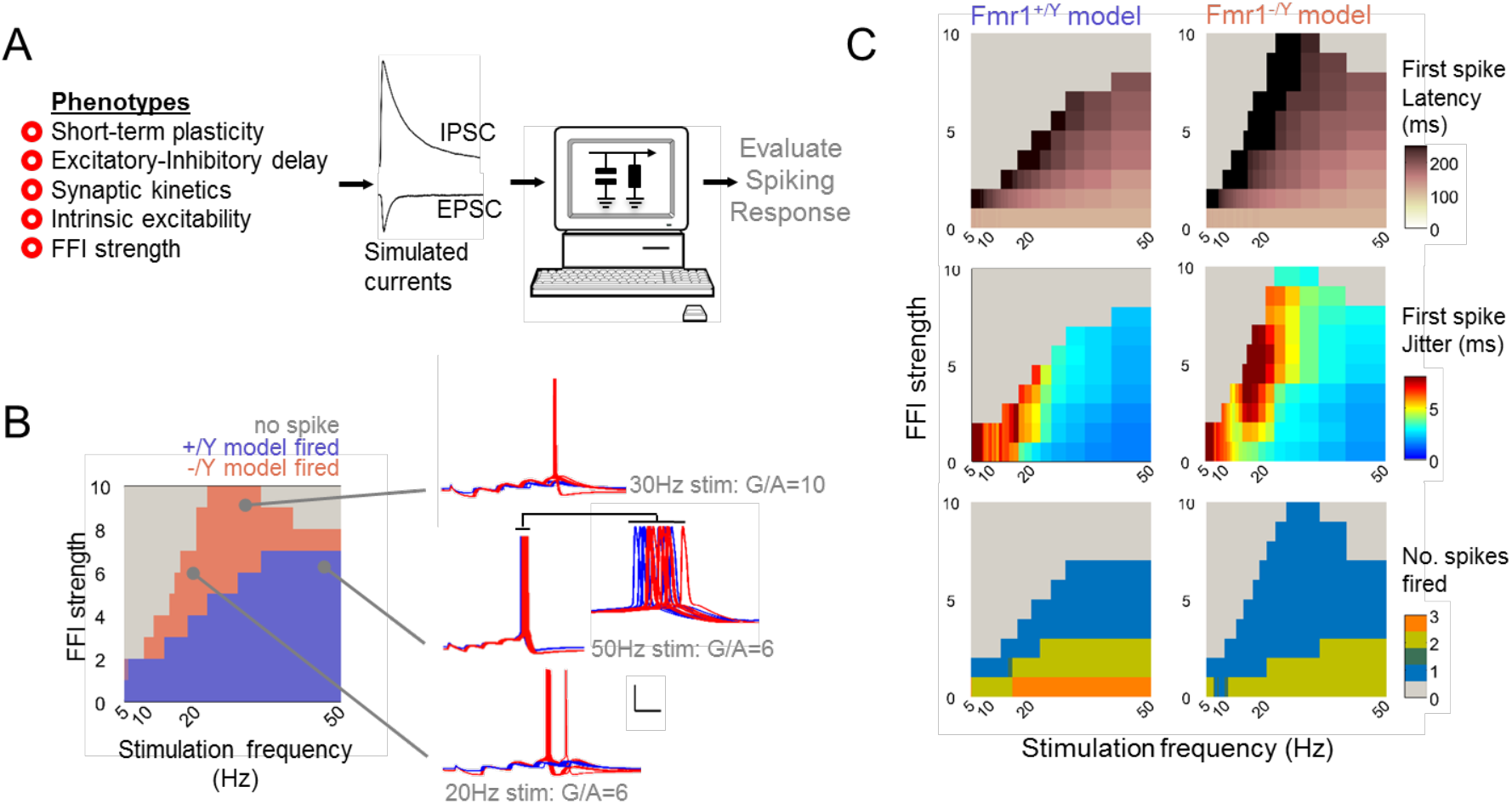
Modelling thalamocortical summation using data-derived parameters recapitulates spiking phenotypes of Layer 4 SCs in *Fmr1-KO* recordings. **A).** Schematic of modelling approach. Left: five grouped covariates measured from WT and KO recordings used in simulation. Centre: Simulated synaptic inputs were tuned with kinetics of recorded currents. Right: Parameter spaces were explored *in silico* to find conditions that either enhanced or suppressed firing in the *Fmr1-KO* model compared to the WT model. **B).** Left: input frequency dependence of simulated spiking responses for model Layer 4 neurons receiving different strengths of FFI (G/A ratios between 0~10 tested). Coloured areas for each modelled genotype indicate combinations of FFI strength/stimulation frequency at which the models fired at least one spike per trial. Note: 1) moderate strength FFI in the WT model prevents spike firing even at high input frequencies 2) the increased number of simulation conditions leading to spiking in the *Fmr1-KO* model, 3) the insensitivity of spiking regulation in the *Frm1-KO* model to inhibitory tone even with FFI strengths elevated to extreme levels (10 trials overlaid). Right: example traces for simulated spiking by the two models at different parameter combinations. Inset: note later and more variable spike times in the *Fmr1-KO* model. Scale: 20mV/50ms. **C).** In addiction to affecting the overall spike firing response shown in B)., genotype dependent effects were observed in the latency, timing variability and count of spikes fired. Spikes fired later and with lower temporally precision in the *Fmr1-KO* model across a broad range of model conditions, even with the FFI strength increased to the extreme values as observed in the *Fmr1-KO* recordings. More conditions led to spiking in the *Fmr1-KO,* despite a slight decrease in numbers of spikes fired per trial across the distribution compared to WT simulations.

We used our single-cell model to explore the role of groups of parameters in regulating spike output generated by thalamocortical input over a range of G/A values (Fig 8). To simplify the model parameter landscape to those involving similar physiological processes, we assigned parameters into four groups. These were namely: short-term plasticity (coefficients constraining synaptic depression behaviour of TC and FFI inputs during repetitive stimulation), excitatory-inhibitory synaptic input delay (lag between simulated TC and FFI input onset times), intrinsic excitability (leak conductances and capacitance of model), and synaptic kinetics (Rise and decay kinetics of synaptic conductance). We then systematically replaced *Fmr1-KO* values with WT values for the various parameter groups to simulate different “rescue” scenarios and measured spike output over a range of G/A values for all of these combinations (four parameter groups to the power of two potential genotypes = 16 simulated scenarios) in response to 5 stimuli at 5-50Hz. For example, when we ‘rescued’ excitatory-inhibitory synaptic input delay on the *Fmr1-KO* background there was an 86% increase in spike output by the model across the parameter space explored, relative to the number of spikes fired by the WT model. Conversely, rescuing intrinsic excitability on the *Fmr1-KO* background produced a 26% reduction in spike output. Thus, manipulation of different parameter groups can have opposing effects on spike output. Overall, by rescuing various combinations of parameter groups we observed that some have antagonistic effects on spike output (e.g. simultaneous rescue of intrinsic excitability and excitatory-inhibitory synaptic input delay) while others drive spike output in the same direction (e.g. excitatory-inhibitory synaptic input delay and short-term plasticity). This analysis indicates that the elevated intrinsic excitability observed in *Fmr1-KO* SCs is a dominating feature driving the changes in spike output. However, changes to other parameters limit the physiological manifestation of this cellular phenotype at the output level of spike generation, suggesting that some of the parameter changes observed in the *Fmr1-KO* thalamocortical phenotype reflect a compensation in response to the underlying pathology that limits circuit dysfunction.

**Figure 8.**
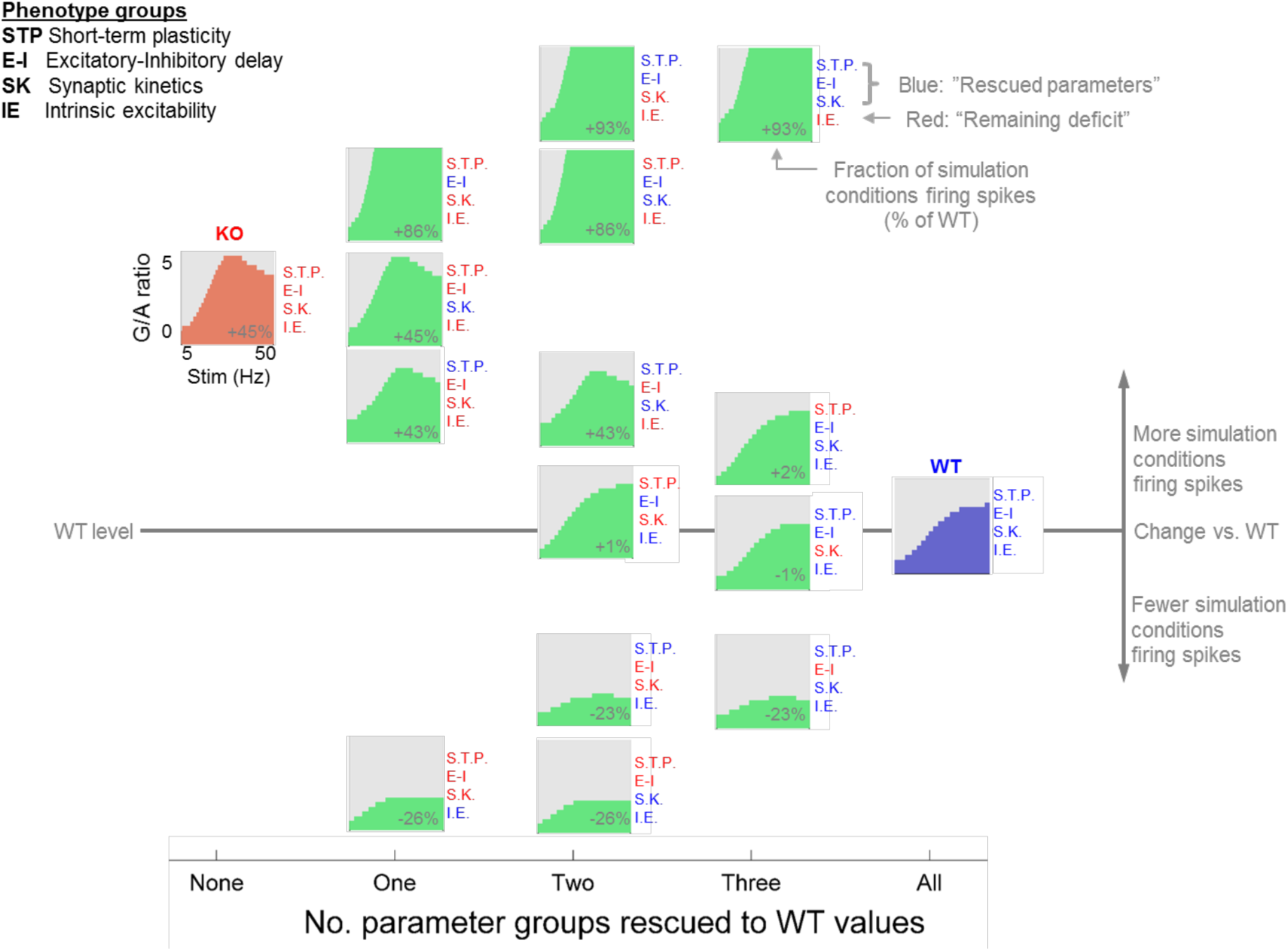
Simulations show the relative contributions of different mechanisms to overall dysfunction in *Fmr1-KO*. Model parameter space explored for 16 different possible combinations of simulated WT and *Fmr1-KO* conditions (4 parameter groups, two possible genotypes, i.e. 4^2^ combinations). Spike firing conditions (in response to five repetitive stimuli) are shown in green for each intermediate rescue scenario as well as *Fmr1-KO* to WT models. For each partial rescue scenario (either with WT or *Fmr1-KO* values, shown in blue and red, respectively), the total count of spike firing conditions across the whole 5-50Hz and 0<G/A<10 range is shown compared to that of the wild-type simulation (i.e. full *Fmr1-KO* model fired >1 spike(s) in 45% more simulated FFI strength/input frequency conditions compared to the WT model).

### Cellular and synaptic deficits in *Fmr1-KO* layer 4 produce network dysfunction

Having explored the mechanisms underpinning the altered transformation of the thalamocortical input in layer 4 *Fmr1-KO*, we next asked how the cellular and synaptic deficits result in altered network activity and what the consequences are for information processing by layer 4. We developed a spiking network model of a cortical layer 4 barrel, based on our electrophysiological recordings (Figures 9A and S6; see also Methods and Table 1). The model consists of a network of interconnected SCs and FS, stimulated by TC input. The great advantage of layer 4 barrel cortex in this regard is the numbers of SCs and FS in a barrel are known as is their probabilistic connectivity (Lefort *et al.*, 2009). The model was randomly connected, constrained by the experimentally determined probabilities (as detailed above) for WT and *Fmr1-KO* phenotypes. Cells were represented as single compartment leaky integrate and fire (LIF) neurons connected by inhibitory (GABA_A_) and excitatory (AMPA and NMDA) synapses displaying short-term plasticity. Additionally, model parameter distributions matched recorded biological diversity in intrinsic neuronal properties, synaptic strength and kinetics, transduction delays and short-term plasticity of synapses (Figure S6). Simulated membrane depolarisation and action potentials were measured in response to the same patterns of train stimulation of TC axons as we used in the slice experiments.

**Figure 9.**
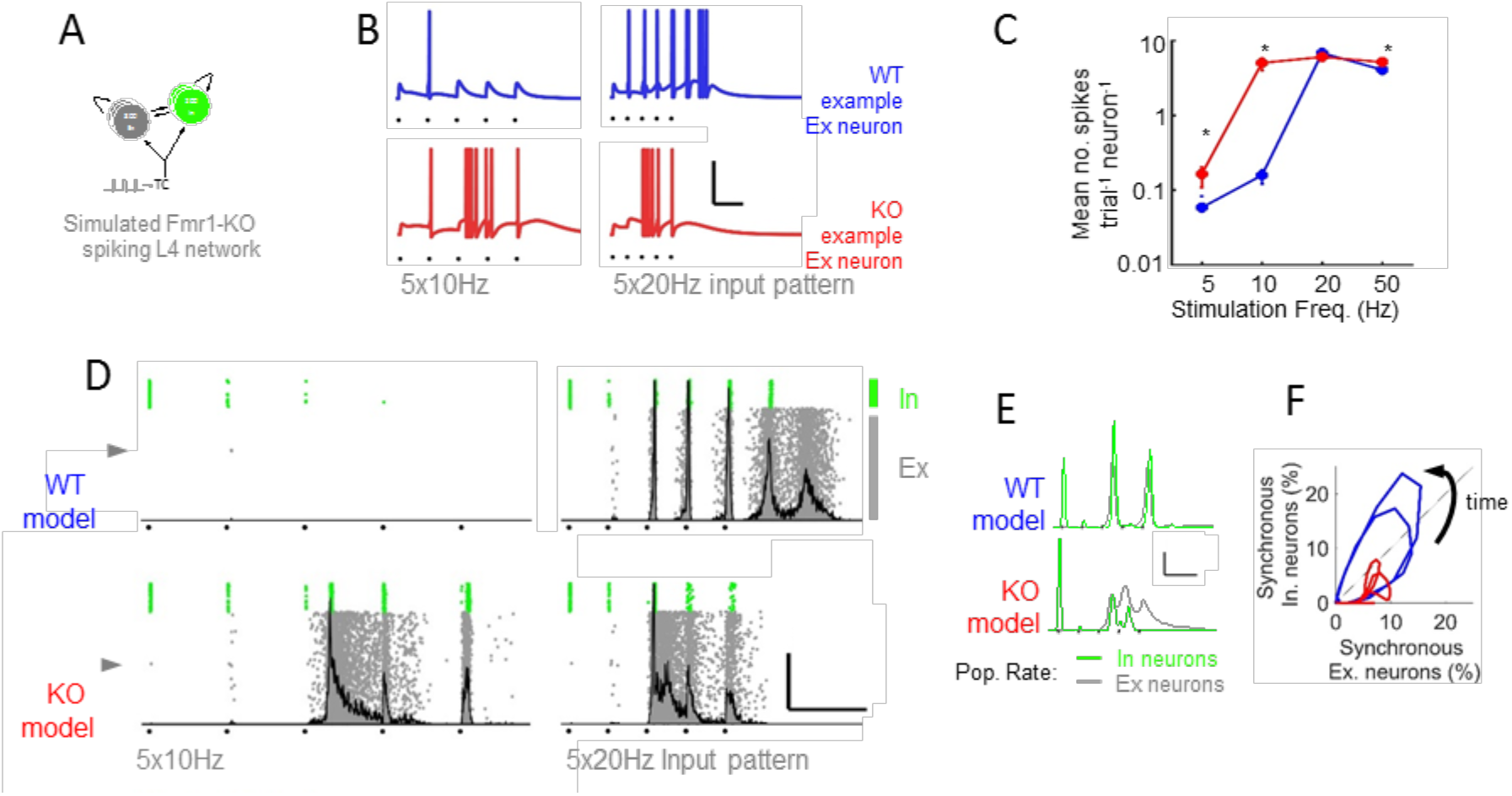
A randomly connected recurrent spiking network model of *Fmr1-KO* layer 4 reproduces pathophysiological features of the population response to TC stimulation. **A).** Schematic of simulated model circuit comprising recurrently connected pools of Ex and In neurons receiving simulated thalamocortical input. **B).** Simulated membrane potential for two example Ex neurons (indicated in D by grey arrowheads) for TC input at 10 and 20Hz stimulation frequency. Black dots indicate stimulus times. Scale: 100ms/50mV. **C).** Frequency-dependent spike output of model Ex neurons. Asterisks denote significantly different spike mean spike counts between models (p<0.05, t-test, 800 neurons per model, 5 random models, average of 10 trials each). **D).** Exemplar full network spike rasters for Ex and In Neurons (grey and green, respectively) showing firing patterns in response to 5x 10-20Hz model thalamocortical stimuli. Population histograms of Ex neurons overlaid in grey. Scale: 100ms, 50% Synchrony **E).** Grand mean Ex (grey) and In (green) population spike density functions (5 random seeds, 10 repeats each) from WT and *Fmr1-KO* simulations. Note impaired E-I population interaction in *Fmr1-KO* simulations. **F).** Phase plot summarizing rhythmic Ex-In population interaction in WT and *Fmr1-KO* models. Note reduced global synchrony and impaired recruitment of Inhibitory neurons in *Fmr1-KO* model.

*Ab initio*, the random model reproduced well the main features of our experimental data in both WT and *Fmr1-KOs*, both at the single cell and at the network level. The *Fmr1-KO* displayed increased subthreshold summation and increased spiking activity to TC input at low train frequencies at the single cell level (Figure 9B, C). Similar to our experimental findings and in agreement with our single-neuron thalamocortical model, the timing of first spikes fired in the *Fmr1-KO* model was delayed and showed decreased inter-trial fidelity (Figure S7A). Similarly, fluctuations in trial-to-trial mean firing rate were enhanced in the *Fmr1-KO* model (Figure S7B). Notably, approximately half of the per-trial averaged numbers of spikes fired by neurons in the recurrently connected model could be accounted for by those fired by our single-cell model lacking recurrent connectivity (Figure 7C cf. Figure S6A) demonstrating a strong thalamocortical contribution to the generation of spiking in layer 4.

At the network level, high frequency TC stimulation evoked a transient reverberating population response, characterised by high rates of simultaneous model neuron firing (highlighted by peaks in the grey firing histograms overlaid in Figure 9D). In the WT model these synchronous population events were initially tightly locked to latter stimuli during the train before transitioning to self-sustaining network activity persisting after the last stimulus. *Fmr1-KO* model network activation required fewer stimuli, was recruited immediately following the first TC-locked spike volley but did not produce the self-sustaining network activity observed in WT. We explored this observation further by examining the stimulus-evoked population firing rates of excitatory and inhibitory neurons in the model network (Figure 9E). The first TC stimulus led to spiking in a greater fraction of *Fmr1-KO* inhibitory neurons compared to WT. Strong reductions in the fraction of firing of inhibitory neurons was observed for successive stimuli for both genotypes, but this was much greater for the *Fmr1-KO* model, consistent with experimental data. In the WT model, firing rate increases in the excitatory and inhibitory populations were strong and temporally matched, leading to a slowly decaying limit cycle (Figure 9F) for recurrent network firing. In contrast, the rates of synchronous firing for excitatory and inhibitory populations were disorganised in the *Fmr1-KO* model, with lower overall synchrony, producing unstable limit cycles.

Overall, the network model captures many of the important features of our experimental data and suggests a mechanism by which single-cell physiological defects in the *Fmr1-KO* layer 4 network (particularly weaker Ex-In connectivity and cellular hyper-excitability) generate impaired network activity.

### Impaired pattern discrimination by a simulated *Fmr1-KO* layer 4 model

Thus far we have demonstrated that a simple data-driven network model of *Fmr1-KO* layer 4 reproduces the main experimentally obtained differences in responses to patterned TC input. Since a major role of barrel cortex layer 4 is the representation of sensory details in the rate and timing of excitatory neurons action potentials (Panzeri *et al.*, 2001), we next examined how individual neurons and small cell ensembles in our WT and *Fmr1-KO* layer 4 models performed when challenged to detect subtle temporal features of simulated sensory input (Figure 10). This was represented as an extra stimulus inserted into the input stimulus train and can be conceptualised as the perturbation of TC neurons’ firing patterns in response to a small surface bump encountered during active sensation.

**Figure 10.**
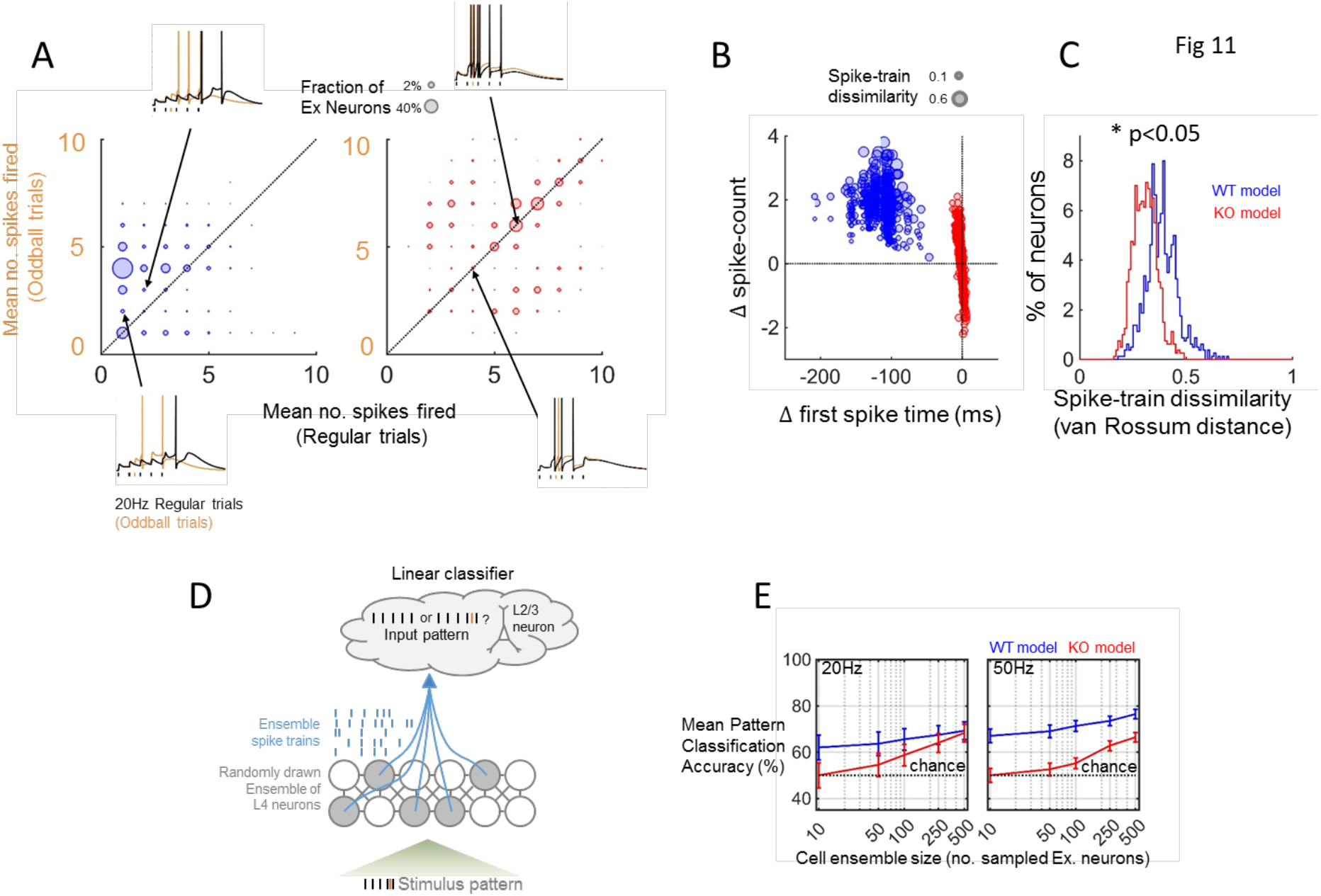
Impaired coding of fine textural detail by neurons and cell ensembles in a recurrent network model of P10-11 *Fmr1-KO* layer 4. **A).** Representation of the presence an extra oddball stimulus inserted into a 20Hz regular stimulus train (orange tick) by changes in spike count fired by Ex neurons. WT model is shown in blue (left), *Fmr1-KO* model is shown in red (right). Circle sizes denote the fraction of neurons in the network represented at each coordinate. Single trial example shown. Insets show simulated membrane potential for example neurons at the locations highlighted in main figure for regular (black) and oddball (orange) trials. Note that while the majority of WT model neurons increased their firing rates for the oddball trial, the majority of *Fmr1-KO* neurons only weakly represented the presence of the oddball stimulus by change in firing rate (bulk of points on diagonal). **B).** Different modes of encoding fine detail of sensory input for simulated WT and *Fmr1-KO* Layer 4 network model spikes. Change in first spike time vs. change in spike rate for Ex neurons in response to additional oddball stimulus. Average of ten trials from 800 Ex neurons across five random network seeds. Size of points denotes mean spike train dissimilarity for individual neurons (van Rossum distance between spike trains on regular and oddball trials). WT neurons typically increased their firing rate and advanced their first spike in response to an extra oddball stimulus. *Fmr1-KO* neurons showed inflexibility in their spike rate but a bidirectional change in first spike time. **C).** Population histogram of van Rossum spike train distances for oddball vs. regular network inputs. Population average oddball sensitivity of individual cells was reduced in the *Fmr1-KO* network model (Kolmogorov-Smirnov test, average of ten trials from 800 Ex neurons across five random network seeds). **D).** Classification of network input from the spike trains of cell ensembles in the Layer 4 model, analogous to readout of input to Layer 4 by Layer 2/3 neurons. Schematic illustration of method for randomly drawing sub-samples of neurons from the Layer 4 network model as inputs to a linear classifier. **E).** Coding of input detail by ensembles of Layer 4 neurons is impaired in *Fmr1-KO* model. Mean leave-one-out decoder cross validation error for WT and *Fmr1-KO* networks for randomly drawn ensembles of varying sizes between 10-500 neurons. Average of 10 random permutations of each ensemble size, four oddball stimulus positions. Simulations were performed on five random network seeds, ten trials for each oddball position and frequency combination. At 20Hz input frequency, ensembles comprised of ten and twenty *Fmr1-KO* neurons performed significantly worse than WT ensembles (t-test, p<0.05), and no better than chance at 20 and 50Hz (one-sample t-test vs 50%, p>0.05) All WT ensemble sizes performed better than chance at both frequencies. At 50Hz, all *Fmr1-KO* ensemble sizes performed worse than corresponding WT ensembles.

The majority of WT Ex neurons detected the oddball stimulus as evidenced by a change in firing rate (indicated by the off-diagonal clustering of points in Figure 10A). However, the majority of simulated *Fmr1-KO* neurons were less likely to detect the oddball stimulus. Additionally, WT neurons represented the presence of an oddball stimulus in the timing of the first spike fired (Figure 10B) such that, despite an interdependence, WT neurons could encode the presence of an oddball as a change in spike rate and/or a shift in first spike timing. Conversely, *Fmr1-KO* neurons showed a reduced change in first spike timing in response to the oddball stimulus. To quantify both of these spike train effects, we used a distance metric that encompasses both spike train and rate (van Rossum, 2001; Victor, 2005) to demonstrate the distribution of single-cell oddball stimulus representations (Figure 10C). The distribution of regular-oddball spike train dissimilarities was uniformly reduced in the KO, implying that the net effect of spike train rate and timing leads to a global reduction across the network in the ability to discriminate a fine temporal feature of simulated sensory input.

The principal output from the layer 4 circuit is a sparse population code provided by the ascending synaptic connection to layer 2/3 neurons (Olshausen and Field, 1997, 2004; Petersen, Panzeri and Diamond, 2001; Golshani *et al.*, 2009b; Jadhav, Wolfe and Feldman, 2009; Crochet *et al.*, 2011), the activity of which plays an instructive role in the development of the layer 2/3 network (Feldman, 2000; Celikel, Szostak and Feldman, 2004). Therefore, it is important to consider how the defects in layer 4 stimulus representation might relay at the subsequent processing level. We therefore used a linear population decoder (Bishop 2011) to classify input stimulus patterns based on the layer 4 output firing pattern (Figure 10D). For population codes formed from ensembles of between 10-500 neurons (i.e. representing convergence of 1/8 ~2/3 of model layer 4 neurons to a layer 2/3 ‘readout’ neuron), more neurons were required for the *Fmr1-KO* model to reach the same degree of classification accuracy as the WT model (Figure 10E). Even drawing 500 neurons (~2/3 of the total excitatory network), population coding of input pattern was impaired in the *Fmr1-KO* model.

Taken together, this result suggests that the developing synaptic pathway from layer 4 to layer 2/3, as schematised here as the layer 4 network’s input pattern classification accuracy would be mis-informative about the quality of sensory input in the *Fmr1-KO* model, and that information gleaned from a larger pool of layer 4 neurons would be necessary for an informative representation of the sensory input.

## Discussion

Altered sensory sensitivities are a common and debilitating feature of FXS and of ASDs more generally. Indeed while the severity of intellectual disability (ID) and epilepsy are highly variable between individual cases of ASD, altered sensory symptoms and/or behavioural responses to sensory information are common to individuals across this spectrum (reviewed in Cascio, 2010; Marco et al., 2011). These sensory abnormalities are distressing for people with ASD and lead to anxiety and social withdrawal, as well as self-injurious behaviour. Unfortunately, we know little about when and how these sensory hypersensitivities arise during development, nor how altered neuronal properties affect experience-dependent development.

We have explored the effects of FMRP deletion on the development of intrinsic and local circuit properties of layer 4 cells in the developing murine somatosensory cortex. In this study, we demonstrate and functionally dissect the intrinsic and synaptic mechanisms that lead to an altered In/Ex balance in layer 4 of somatosensory cortex immediately prior to the onset of active sensory exploration but after the critical period for synaptic plasticity at TC synapses. Surprisingly, we show that FMRP deletion results in an increase in In/Ex ratio in some layer 4 SCs. However, this increase in inhibition is rapidly lost at physiological stimulation frequencies as a result of disruptions in synaptic kinetics and short-term dynamics of feed-forward inhibition. The distributed nature of the physiological changes that result from the loss of FMRP indicate that compensatory changes may be activated even at these early stages of development. Indeed, our modelling indicates that many of the changes observed in the *Fmr1* KO move the network response in opposite directions and thus are antagonistic to one another in regulating cell excitability. Furthermore, modelling these phenotypes predicts an altered temporal coincidence detection of thalamocortical inputs at a range of physiologically relevant stimulation frequencies, a prediction that is borne out by experimental data. Together these data indicate that the sensory information being relayed to layer 3 is dramatically altered at this stage of development, which would likely have a deleterious effect on experience-dependent development and hence sensory processing at later ages. They also highlight the power of integrating experimental and mathematical approaches to understanding and forming testable hypothesis about the nature of the developmental mechanisms underlying neuronal circuit dysfunction in ASD.

### Distributed nature of the physiological phenotypes associated with the loss of FMRP

Genetic deletion of FMRP causes a range of cellular phenotypes that could be causally linked to the circuit hyperexcitability and sensory hypersensitivities (Contractor, Klyachko and Portera-Cailliau, 2015). However, which phenotypes arise as a direct loss of FMRP as opposed to homeostatic/compensatory changes that regulate cellular and/or circuit excitability is not known. In this context, many of the cellular phenotypes described here have opposite effects on excitability on circuit output, strongly suggesting that some, if not all, are likely compensatory changes driven downstream of the direct effects of FMRP deletion. That said, each of these phenotypes could provide a potential for therapeutic intervention and have a distinct contribution to the overall hyperexcitability phenotype.

### Inhibition/Excitation balance does not predict cellular excitability

An altered In/Ex balance has been causally linked to a range of psychiatric disorders from ASD to schizophrenia (Rubenstein and Merzenich, 2003; Chao *et al.*, 2010; Yizhar *et al.*, 2011; Lisman, 2012; Uhlhaas, 2013). In primary somatosensory cortex, the In/Ex balance is essential for regulating both the frequency (spike rate) and quality (spike timing) of action potentials, and hence for accurately processing sensory input (Panzeri *et al.*, 2001). In particular, the precise temporal tuning of excitation and inhibition is critical to signal processing of cortical networks (Haider and Mccormick, 2009). Here, we find an increase in the In/Ex balance in a subset of layer 4 SCs that causes an overall increase in the In/Ex at the population level despite the observation of an ectopic pool of *Fmr1-KO* SCs lacking functional FFI. However, this elevation in In/Ex balance is accompanied by a paradoxical gain in overall excitability because of elevated SC intrinsic excitability and an increase in the delay between TC excitation and feed-forward inhibition onto spiny stellate neurons (Ex-In lag). This delay results in a broadening of the TC EPSP and a large increase in the summation of EPSCs following TC stimulation that is especially pronounced at higher stimulation frequencies. Hence, the relative increase in inhibitory tone is less effective at truncating the incoming excitation due to its late arrival. Compounding this increase in excitatory summation, we find a more dramatic loss of inhibitory relative to excitatory tone during physiologically-relevant repetitive stimulations. The rapid decrease of inhibition results in a further increase in summation of the EPSPs, especially during higher frequency stimulations, where repetitive TC input arrival occurs faster than the postsynaptic voltage relaxation due to the intrinsic and synaptic decay kinetics of the SCs and their inputs. Thus, although initial assessment revealed an increase In/Ex balance for layer 4 SCs, alterations in the synaptic kinetics and short-term plasticity result in an increase in cellular excitability leading to a loss of pattern discrimination. These data highlight the vagaries of simply examining In/Ex imbalance as a key characteristic of circuit dysfunction. It is important to remember that the In/Ex balance is dynamically regulated (Hasenstaub *et al.*, 2005; Haider and Mccormick, 2009; Gentet *et al.*, 2010; Crochet *et al.*, 2011; Haider, Häusser and Carandini, 2012) and is only one factor that regulates neuronal processing within a complex circuit and its role in mediating sensory dysfunction (O’Donnell et al. 2017; Golshani et al. 2009).

### Abnormal intrinsic neuronal membrane properties

Our finding of an increase in input resistance of layer 4 SCs is in good agreement with previous findings for this cell type (Gibson *et al.*, 2008) and is perhaps the most consistent physiological feature across neuronal cell types in the *Fmr1-KO* mouse (Gibson *et al.*, 2008; Meredith and Mansvelder, 2010; Olmos-Serrano *et al.*, 2010; Brager, Akhavan and Johnston, 2012). This elevation in input resistance and resultant slowing in membrane time constant, raises the impedance to sinusoidal stimuli. By increasing the intrinsic sensitivity of *Fmr1-KO* SC neurons to low frequency stimulation this has the compound effect of many more action potentials being generated in response to low frequency stimuli, and a dampening of responsivity to high frequency inputs. Importantly, these input resistance-driven changes in intrinsic properties also result in a phase shift in the voltage response to input currents, for example producing a phase shift in action potential generate. Such a shift in phase can have dramatic effects on spike-time dependent plasticity (STDP, Minlebaev et al. 2011) which may contribute to the shift in the timing of the critical period for LTP at thalamocortical synapses (Harlow et al., 2010). Crucially, the maturation of the layer 4 excitatory network in the brief developmental window beyond P10 is sensitive to sensory experience (Ashby and Isaac, 2011) and relies on spike timing-dependent plasticity mechanisms (Egger, Feldmeyer and Sakmann, 1999). The cellular and circuit disruptions to spike time representation reported here would be expected to dramatically alter experience-dependent development of primary somatosensory cortex in *Fmr1*-KO mice and could contribute to the circuit hyperexcitability reported at older ages. Consistent with this idea, impaired STDP is reported in the *Fmr1-KO* mouse prefrontal cortex (Meredith, Holmgren and Weidum, 2007).

Our findings indicate that the circuit deficits in layer 4 of somatosensory cortex arise in *Fmr1* KO mice arise from a range of cellular deficits however, their relative contribution to the overall layer 4 circuit function and hence the information propagated to layer 2/3 in not known, prompting our use of circuit modelling.

### Modelling local circuit dysfunction in Fmr1-KO mice

The approach taken in this study was to build a mechanistic understanding of a cellular and circuit dysfunction through combined and bidirectional physiology and predictive modelling. By studying seemingly small pathophysiological changes in the component parts of a well-described dynamic circuit, our models were able to predict large physiological increases in the sensitivity to ethological stimuli. The single cell FFI model specifically predicted the altered summation properties at a range of stimulation frequencies that were verified through experimentation. Furthermore, by substituting each parameter or combination of parameters from *Fmr1-KO* animals with that from their WT counterparts, we were able to begin examining the relative contribution of each phenotype to the overall local circuit dysfunction.

Taken as a snapshot in time, the cellular model does not distinguish between the phenotypes that result directly from the loss of FMRP and which arise secondarily to compensate for these changes. However, it does provide a powerful framework to study compensation within neocortical neurons that arise from particular genetic mutations. It can also be used to examine the effects of targeted treatments, for example by simulating the effect of a single parameter repair or combination repair strategy, as we do in the current study. This has the potential to reduce cost, time and animal usage. The model demonstrates how many of the changes observed in the *Fmr1-KO* move the network activity in opposite directions and thus are likely compensatory to one another. These compensatory changes likely reflect mechanisms by which the impact on overall network function is minimized.

To further address the question of causal vs. compensatory physiological disruptions, three main approaches will be necessary, firstly through physiological investigations spanning development timepoints (this study, Gibson et al. 2008; Harlow et al. 2010) to evaluate the developmental evolution of features of physiological dysfunction, possibly in combination with chronic sensory manipulation (e.g. whisker deprivation) or carefully timed transient rescue interventions. This approach can be augmented by (secondly) analytical mathematical methods and (thirdly) numerical simulations. The derivation of exact analytical expressions governing network activity is notably challenging in the face of the complex, non-linear circuit dynamics (Transtrum, MacHta and Sethna, 2010; O’Leary, Sutton and Marder, 2015) but in reduced circuit or parametric statistical models (e.g. Panas et al. 2015) offers powerful insight into which parameter(s) are dominant and which the network is robust to upon manipulation (highlighting stiff vs. sloppy parameters, Panas et al. 2015; Machta et al. 2013). Numerical simulation offers a complementary approach to studying network homeostasis and development (Leary *et al.*, 2009; Litwin-Kumar and Doiron, 2014). These crucially provide predictions for the stochastic evolution of network structure and dynamics under different combinations of interventions, thereby comparing simulated network developmental trajectories from a common starting point. We introduce here a machine learning approach to classifying stimulus identity from multi-neuronal firing patterns. This could be extended to probe the evolving information capacity of model networks at different stages of *in silico* development. One promising approach is to compare a (model) circuit’s observed information capacity to that of a theoretically optimal model, in which a trade-off has been met for factors such as neural response correlation and response redundancy (Tkacik *et al.*, 2010). The derivation of such an optimal circuit model could be hampered however by the adoption of animals’ alternative behavioural strategies to extract relevant information from sensory input, as has been reported in Autistic individuals (Happé, Frith et al., 2009; Livingston and Happé, 2017)

Future experiments combining experimental manipulations of activity levels with model predictions will be able to elucidate which cellular parameters remain malleable and hence what pharmacological interventions may be more effective in restoring circuit function. These predictions can subsequently undergo rigorous testing by physiological experimentation. Such an approach would also be a valuable tool for examining the convergence of pathophysiology in disease mechanisms affecting circuit function across a range of genetic models.

While the cellular model is very useful for analysing the role of particular cellular and synaptic phenotypes in regulating neuronal integration and output properties, it does not encompass the recurrent network activity initiated by thalamocortical activity. Therefore, we generated a network model to investigate the nature of the output of the of the layer 4 circuit and hence the activity that will subsequently influence the development of cortical circuits further down the sensory processing hierarchy and hence the altered circuit excitability believed to underpin sensory hypersensitivity (Gibson *et al.*, 2008; Huang *et al.*, 2011; Gonçalves, Anstey, Golshani and Portera-cailliau, 2013). The primary goal of this endeavour was to the understand the quality and fidelity of the information leaving layer 4 in *Fmr1-KO* compared to WT which is key to driving experience-dependent development of layer 2/3 neurons (Feldman, 2000). Using data-tuned cell models as a starting point, we incorporated experimentally-derived connection probabilities with known layer SC and FS cell numbers and GABA, NMDA and AMPA currents. The model predicts a decrease in spike firing and loss of trial to trial fidelity in the *Fmr1-KO* animal, accurately reflecting our data obtained using patch-clamp recordings from slices. To simulate a subtle temporal sensory disturbance during active whisking, we examined the effect of an oddball stimulus introduced during the stimulus train. The WT model was able to detect the presence of an oddball stimulus through a change in the firing rate and timing of the spikes fired by Excitatory neurons. On the contrary, the *Fmr1-KO* model was less likely to alter its firing rate and the timing of the first spike in response to the oddball stimulus, indicating a robust decrease in the circuit’s ability to discriminate changes in the sensory environment. These findings strongly suggest that the ability of the layer 4 circuit to relay details about patterned sensory information, a key feature of experience-dependent plasticity, is dramatically reduced in *Fmr1-KO* mice. The consequence of reduced sensory driven plasticity would be a retardation of circuit development. Importantly, Bureau and colleagues (2008) found a delay in the developmental increase in connection probability between layer 4 to layer 2/3 cells in somatosensory cortex of *Fmr1-KO* mice that was mirrored by a delay in the pruning of SC axons projecting to layer 2/3. This delay could be accounted for the shift in the sensitive period for inducing LTP at TC inputs in layer 4 in somatosensory cortex which is delayed in *Fmr1-KO* animals (Harlow *et al.*, 2010). Findings from our model now show that the quality of the information being transmitted by the layer 4 to layer 2/3 cells is dramatically reduced. Importantly while some key features of somatosensory cortex development are delayed, this is not a global phenomenon. For example, layer 4 SCs show typical morphological development, with a typical pattern of spinogenesis, synaptogenesis more generally is unaltered for these cells (Till et al. 2012; Harlow et al. 2010 and our subsequent unpublished data). They even show the typical activity-dependent restriction of their dendrites to the regions of thalamocortical axon terminals (Till *et al.*, 2012). The findings provide support for the idea that a mismatch in the maturation of physiological and/or anatomical properties that are necessary for the subsequent development of circuit function (Meredith et al. 2012).

This mismatch in developmental trajectories may subsequently induce altered development in downstream areas. Antoine et al. (2018) recently reported a decrease in In/Ex balance in layer 2/3 of barrel cortex in *Fmr1-KO* mice. However, this change in the In/Ex balance did not result in an increase in cellular excitability but instead acted to stabilise synaptically-driven output of the neuron, strongly suggesting that neurons alter their In/Ex balance to compensate for other changes in the neuron’s physiological properties. Intriguingly, Gainey et al. (2018) found a similar compensatory change in In/Ex following experience-induced plasticity to whisker trimming, suggesting that some features of excitatory and inhibitory circuit plasticity that underlie this effect (House *et al.*, 2011) are spared in the *Fmr1-KO*. These findings are particularly interesting in terms of our data from P10-P11 animals. At P10, the changes In/Ex balance reported here do not normalise the output of the neuron possibly because of the early developmental stage studied. Instead, we find a dramatic reduction in the information being propagated to layer 2/3 at this age. Hence, for a layer 2/3 neuron, the loss of FMRP essentially mimics sensory deprivation, which could explain the changes in In/Ex balance seen in layer 2/3 at P21 observed by Antoine et al. (2018). Compensatory changes in the In/Ex balance were also seen in other models of ASD (Antoine *et al.*, 2018). Together these findings suggest that altered neuronal physiology early in development drive altered sensory processing and homeostatic changes in In/Ex balance that stabilise neuronal firing.

## Materials and Methods

### Mice

*Fmr1-KO* mice (B6.129-Fmr1^tm1Cgr^, RRID:MGI_MGI:3815018) were obtained from the Contactor laboratory (Northwestern University, Chicago IL, USA). Mice were maintained on a C57Bl6/J background. Hemizygous male *Fmr1-KO* (*Fmr1^-/Y^*) and wild-type littermate (WT, *Fmr1^+/Y^*) pups were used at postnatal days 10-11 (P10-11). All procedures were carried out according to UK Home Office and NIH IACUC guidelines for animal welfare. N’s of neurons/animals for each experiment are indicated in the individual figure legends.

### Brain slice preparation

Mice were decapitated and brains were rapidly removed and placed in ice-cold carbonated (95% O_2_ / 5% CO_2_) cutting solution. Thalamocortical brain slices (400μm thick) were prepared according to Agmon and Connors (1991) using a vibrating microslicer (Leica VT1200). Slices were transferred to artificial cerebro-spinal fluid (aCSF) and stored at room temperature. aCSF contained (in mM): NaCl 119, KCl 2.5, NaH_2_PO_4_ 1, NaHCO_3_ 26.5, Glucose 11, MgSO_4_ 1.3, CaCl_2_ 2.5. Cutting solution was identical except for substitution with 7mM MgSO_4_ and 0.5mM CaCl_2_.

### Electrophysiology

Patch-clamp recordings were performed using Cs^+^-based internal solution containing (in millimolar): CsMeSO_4_ 130, NaCl 10, HEPES 5, EGTA 0.5, Mg-ATP 4, Na-GTP 0.3, pH 7.4 at 32°C, adjusted to 285mOsm. Junction potential was uncorrected. Slices were superfused with carbogenated aCSF at 32±0.5°C at 8ml/min. Patch recordings were performed with Multiclamp 700B amplifiers (Molecular Devices), using pipettes (4-7MΩ open bath resistance) fabricated from thin-walled borosilicate glass (Warner Instruments). Signals were low-pass Bessel filtered at 10KHz and digitized at 20KHz using either a Molecular Devices Digidata 1322A board and pClamp 10.2 for recordings performed in the USA, or a National Instruments PCI-6110 board, using acquisition software custom-written in C++/MATLAB incorporating modules from Ephus (http://www.ephus.org) (Suter, 2010) for recordings performed in the UK. Custom stimulating electrodes were manufactured using twisted Ni:Cr wire. For TC fibre stimulation, stimulation electrodes were inserted into VB thalamus. Biphasic, 100μs constant voltage pulses were delivered by an optically-isolated stimulus generator (AMPI systems) at 0.03Hz inter-pulse frequency. Layer 4 SCs were recorded in barrels that showed >100μV TC-evoked field potentials. Cells were selected for recording using DIC optics under infrared illumination based upon somatic morphology and laminar position. Resting potential was measured in bridge balance configuration immediately after breaking in. Recording quality was monitored on-line incorporating the following criteria: resting potentials more hyperpolarized than -50mV, stable (<25MΩ, <20% drift) access resistance and holding currents <100pA. Cell-attached recordings were performed according to Perkins (2006). Briefly, stable greater than gigaohm seals were obtained between electrode and neuron with minimal suction to minimise mechanical membrane stress. Recordings were performed in voltage-clamp configuration by holding cells at pipette potentials that were empirically determined in current clamp to minimise holding current. Recordings were accepted if all action currents (“spikes”) fired were <-200pA in peak amplitude, seal resistance and holding current were stable for the duration of the experiment. In our hands, the statistics of network-evoked firing responses were stable for over an hour (Figures 1, 3 and data not shown), in contrast to Alcami *et al.* (2012).

### Impedance profiling

Sinusoidal current waveforms (“ZAPs”), with *I_inject_*(*t*) = *A* sin (2*πft*) were injected into current-clamped neurons, with amplitude (*A*) of ±40pA and frequency (*f*) increasing between 0.5-50Hz over 25s. Fourier-transformed voltage and current waveforms were used to derive the neurons’ complex impedance *Z̅*(*f*) according 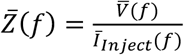 (Cole and Curtis, 1936, 1938; Puil, Gimbarzevsky and Miura, 1986; Carandini *et al.*, 1996), where superscripts denote the Fourier operator (Koch, 1999). The impedance magnitude, | *Z̅*(*f*)| is thus a real valued number between R_membrane_ for sustained DC current (i.e. Ohmic resistance, *Z̅*(*f* = 0) = R_membrane_) and zero, decreasing to zero in the limit of *f* → ∞. The phase shift *ϕ*_(*f*)_ between the input current and voltage output was taken as the inverse tangent of the ratio between the real and imaginary components of the complex impedance:

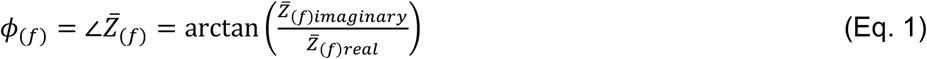

For Bode filter analysis of impedance profiles, the frequency-dependent system gain is expressed as log-power ratios of impedance at higher frequencies relative to that at the lowest tested (f=0.5Hz). Little additional output attenuation compared to DC is expected for either genotype at 0.5Hz, i.e. where *R_membrane_* ≈ *Z*_(*f* =0.5*Hz*)_, hence this is a fair normalization strategy (Zemankovics *et al.*, 2011). Model first-order RC low-pass filter responses were assumed as:

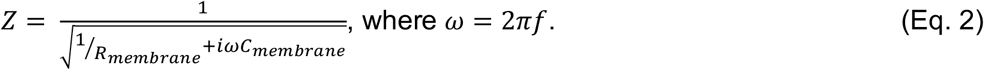

The bandwidth, or “cut-off” frequency *f_cut–off_*, of an ideal first-order low-pass filter, above which the output power relative to DC is attenuated by greater than -3dB (i.e. approximately halved) is given as:

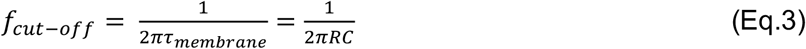

Where here R and C are R_membrane_ and C_membrane_, respectively. Above *f_cut–off_* the voltage response decays with a frequency-dependent roll-off of -20dB/decade. It can therefore be expected that an increase in either R_membrane_ and C_membrane_ will, by increasing the membrane time constant, lower the cut-off frequency of the filter effect.

### Oscillatory spike locking

To find firing threshold, current clamped neurons were first depolarized by bias current injection until repetitive action potential firing was observed. Holding potentials were then corrected to 5mV hyperpolarized from this value at the start of each trial (“V_Baseline_”, Lawrence et al., 2006). Sinusoidal current stimulation was repeated five times for each frequency. To avoid affecting AP initiation by fluctuating voltage level, cells in which V_Baseline_ changed by >2mV were discarded. Phase-locking of AP firing was assessed by registering each AP’s peak time to the phase of the injected current waveform for each cycle, calculating using the Hilbert transform in MATLAB.

### Data analysis

All data were analysed using custom-written routines in MATLAB, except for additional toolboxes detailed below. Where beneficial to improve analysis speed, C++ code was compiled and called from within MATLAB as MEX files.

### Statistical analysis

Data is shown summarised as mean±S.E.M. unless otherwise stated. Individual parameter distributions were tested for normality (Kolmogorov-Smirnov tests, p>0.05) before comparisons were made. Non-parametric tests were used under conditions of deviation from normality, as stated in the individual figure legends.

### Spike train statistics

For spike probability analysis, spike times from cell-attached recordings were down-sampled to 1ms resolution and converted to spike probability functions. In this approach each spike time is discretized at 1ms and convolved with a Gaussian kernel with a standard deviation of 5ms. Each spike’s resulting total integral was adjusted to 1 across a range of 3 standard deviations. Correspondingly, spike times are smoothed such that a spike occurs at time *t* contributes an extra ~0.1 spike probability to that of the next spike if it arises 30ms later. The resulting spike density convolution is a continuous function rather than a discrete train; more amenable to comparison between trials and recordings, and less sensitive to binning artefacts than with a raw spike count histogram approach.

### Thalamocortical integration simulations

A single compartment model neuron consisting of a soma with one dendrite was instantiated in NEURON (Hines *et al.*, 2002). Dendrite length and global leak conductance were tuned to match genotype mean values for input resistance and whole-cell capacitance of *Fmr1-KO* and wild-type recordings. To recreate TC and FFI inputs, two “Exp2Syn” model synapses were placed at the soma, with reversal potentials of 0mV and -71mV, respectively. Synaptic kinetics (rise and fall times) matched voltage clamp data. The model TC input was activated with a peak conductance of 1nS, yielding an ~60pA peak inward current from a leak reversal potential of -60mV. The model FFI input was then activated with an onset latency matching voltage clamp recordings. To recreate the effects of FFI under conditions of different G/A ratios, peak conductance that was varied on successive trials relative to the peak TC conductance between 0-10x, in increments of 0.5.

### Short-term plasticity model

Short-term depression dynamics of TC-EPSC and FF-IPSC current components were quantified using bi-exponential decay fits to trial-averaged voltage clamp recordings, normalized to initial steady-state current amplitudes. Curves in Figure 5c are best fits to within-cell measurements of G/A ratios, normalized to steady-state G/A ratio for each cell. Least-squares optimal fits to G/A ratio depression were obtained by using the “fminsearch” algorithm in MATLAB to obtain terms for underlying TC-EPSC and FF-IPSC components. For implementation in NEURON (v7.3), Depressing current inputs were re-fit with a phenomenological model of STP (Varela *et al.*, 1997). Briefly, starting from steady-state amplitude *A_0_*, the stimulus evoked response amplitude *A* of each current was multiplied by two dynamic depression factors, *D_1_* and *D_2_* (constrained <1):

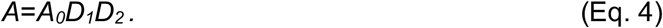

After each stimulus, *D_1_* and *D_2_* were multiplied by constants *d_1_* and *d_2_*, representing the amount of depression per presynaptic action potential (i.e. *D*_1_ → *D*_1_ *d*_1_, *and D*_2_ → *D*_2_ *d*_2_). Between stimuli, *D* variables recovered exponentially back towards 1 with first–order kinetics governed by recovery time constants *τ_D1_* and *τ_D2_*:

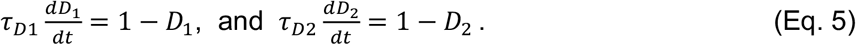

Accumulation of synaptic depression was therefore observed when the inter-stimulus interval was shorter than time required for recovery. Synaptic rise and decay time constants, as well as synaptic plasticity constants *d_1_, d_2_, τ_D1_* and *τ_D2_* were obtained from the genotype mean synaptic kinetics and short-term depression of EPSCs and FF-IPSCs individually. Fitting of short-term depression terms was performed on normalized synaptic depression curves for 5,10, 20 and 50Hz TC stimulation, equally weighted and optimised using a Levenberg-Marquardt search implemented in MATLAB.

### Layer 4 network simulations

A recurrently connected spiking network of single-compartment conductance-based Leaky Integrate and Fire (LIF) neurons was instantiated in MATLAB. The sub-threshold potential of the *i^th^* neuron evolved according to:

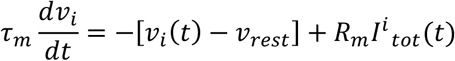

where *I^i^ tot*(*t*) is the summed synaptic current from recurrent and external inputs at time *t*. Spikes were emitted at *t^f^* = {*t*|*v*(*t*) = *v_threhsold_*} when *v* > *v_threhsold_* leading to the reset condition *v*(*t* + *dt*) → *v_reset_* < *v_threhsold_* for a finite refractory period of 1.5ms.

*I^i^ tot*(*t*) was calculated at each time step as:

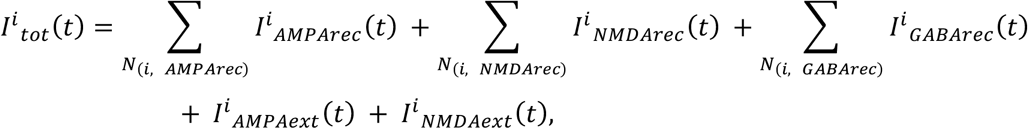

with *N*_(*i, AMPArec*)_, *N*_(*i, NMDArec*)_ and *N*_(*i, GABArec*)_ being the set of *N* recurrent synapses projecting onto the *i^th^* neuron, and *I^i^* _*AMPArec*_ (*t*), *I^i^* _*NMDArec*_ (*t*), *I^i^ _GABArec_* (*t*), *I^i^* _*AMPAext*_ (*t*), *I^i^* _*NMDAext*_ (*t*) the synaptic inputs from recurrent AMPA, NMDA and GABA and external AMPA and NMDA synapses, respectively.

Synaptic input current depended also on the postsynaptic membrane potential:

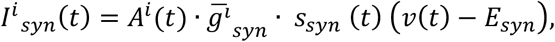

where *A^i^*(*t*) is the short-term depression state of synapse *i* described by Eq. 4, *ḡ_syn_* and *E_syn_* are respectively the maximal synaptic conductance and reversal potential of the synapse. *s_syn_* (*t*) is a delayed-sum-of-exponentials (“alpha”) synapse model (after Dayan & Abbott 2000):

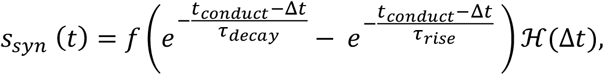

in which *t_conduct_* is a finite synaptic conduction delay, Δ*t* > −∞ is the elapsed time *t* – *t^f^* since the time of the last presynaptic spike. The Heaviside step function ℋ ensures causality in time. The function *f* is an amplitude normalisation factor:

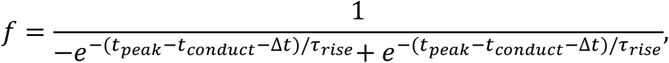

where the amplitude of the conductance is maximal at time *t_peak_*:

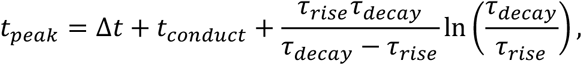

NMDA conductances displayed additional voltage dependence:

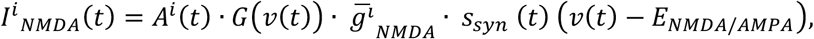

In which *G*(*v*(*t*)) describes the voltage-dependent Mg^2+^-sensitive blockade of NMDA receptors:

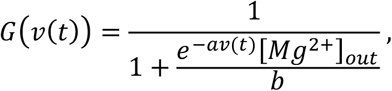

with *a* =0.062mV^-1^ and *b* = 3.57mM (Jahr and Stevens 1999, Gabbiani and Koch 1994) and using our experimental extracellular Magnesium concentration [*Mg*^2+^]_*out*_ of 1.3mM.

### Network architecture

800 Excitatory and 150 Inhibitory neurons were randomly connected with probabilities obtained experimentally in Figure 2, excluding autapses. All classes of synaptic connection were randomly assigned fractional weights up to *ḡ_syn_*, with the exception of Ex-Ex connections, which were log-normally distributed in line with our experimental findings (Figure 3E) and previous reports (Petersen and Sakmann, 2000; Lefort *et al.*, 2009; Ashby and Isaac, 2011). Simulated external thalamocortical inputs was provided to 80% of Ex and In neurons. *ḡ_syn_* of external NMDA and AMPA were jittered between trials and thalamorecipient neurons were shuffled to introduce variability between trials. Short –term plasticity at TC synapses was modelled using the same fits to data as described above. Simulations (1000ms) were carried out using forward Euler integration at 0.5ms resolution. Spike times were discretized to 1ms resolution. Modelling results show responses of network simulation to N=10 trials for each stimulation condition each from 5 random network seeds.

### Parameter distributions

Cell-intrinsic, synaptic and connectivity parameters used are detailed in Supplemental Table 1. Where two numbers are shown, they represent mean and standard deviation of jittered variables. All parameters were assumed to vary independently, excluding input resistance and membrane capacitance, which displayed both private and shared variance.

### Spike train metrics and ensemble decoding

Spike train distance metrics were computed with the method described by van Rossum (2001) with an decay time constant of 50ms. Ensemble decoding was performed using a multivariate linear decoder using ensemble spike trains as covariates. To prevent over-fitting, pooled covariance matrices for regular and oddball conditions were regularised by a factor of 0.05 on the identity matrix. Decoder accuracy was evaluated with ten-fold leave-one-out cross-validation, with decoders trained on nine trials asked to predict the stimulus condition that lead to the outcome of the remaining withheld trial. Training/test sets were split 70/30% and independent evaluation sets were used.

### Code/model availability

NEURON code for the single cell model and MATLAB/C++ code for the Layer 4 network model will be made publicly available on ModelDB and Github. MATLAB/C++ code for analysis of data and simulations will be available from the authors upon reasonable request.

